# Sequencing of *N*^6^-methyl-deoxyadenosine at single-base resolution across the mammalian genome

**DOI:** 10.1101/2023.01.16.524325

**Authors:** Xinran Feng, Xiaolong Cui, Li-Sheng Zhang, Chang Ye, Pingluan Wang, Yuhao Zhong, Zhong Zheng, Chuan He

## Abstract

While DNA *N*^6^-methyl-deoxyadenosine (6mA) is abundant in bacteria and protists, its presence and function in mammalian genomes have been less clear. We present Direct-Read 6mA sequencing (DR-6mA-seq), an antibody-independent method to measure 6mA at base-resolution with high sensitivity. DR-6mA-seq employs a unique mutation-based strategy to reveal 6mA sites as misincorporation signatures without any chemical or enzymatic modulation of 6mA. We validated DR-6mA-seq through successful mapping of the well-characterized G(6mA)TC motif in the *E. coli* DNA and identified 6mA sites in the mammalian mitochondrial DNA. As expected, when applying DR-6mA-seq to mammalian systems, we found that genomic DNA (gDNA) 6mA abundance is in general low in most mammalian tissues and cells; however, we did observe distinct gDNA 6mA sites in mouse testis and glioblastoma cells. DR-6mA-seq provides an enabling tool to detect 6mA at single-base resolution with high sensitivity for a comprehensive understanding of DNA 6mA in eukaryotes.

## Introduction

Covalent DNA modifications can be crucial epigenetic marks in diverse biological systems.^1^ Among them, *N^6^*-methyl-deoxyadenosine (6mA) is a prevalent modification in the genomes of bacteria and protists, playing critical roles in the restriction modification system, DNA repair, replication, transcription, nucleoid segregation, and gene expression regulation.^2^ Albeit well-characterized in prokaryotes and lower eukaryotes, 6mA is considered a rare DNA modification in the genomes of high eukaryotes, especially mammals.^3^ The presence and functional roles of 6mA in the mammalian genome have not been clear largely due to the lack of sensitive and accurate methods to detect this modification at the genome-wide scale.

Given the considerably low modification levels of 6mA in most mammalian genomes, highly sensitive and accurate sequencing methods are required in order to detect the presence of 6mA.^3, 4^ Ultra-high performance liquid chromatography coupled with mass spectrometry (UHPLC-MS/MS) has been widely used to quantify global 6mA levels, but the readout is extremely vulnerable to bacterial DNA contaminants arising from cell culture infection, plasmids, and reagents.^4, 5^ Meanwhile, the 6mA concentrations in mammalian genomic DNA (gDNA) samples are usually around the detection limit of UHPLC-MS/MS, further exacerbating effects from bacterial DNA contamination.^6, 7^ These factors highlight a crucial need to develop high-sensitivity, base-resolution, and whole-genome mapping methods for 6mA sequencing. DNA immunoprecipitation sequencing (DIP-seq), sometimes coupled with exonuclease digestion (6mA crosslinking exonuclease sequencing; 6mACE-seq or ChIP-exo), has been extensively utilized to generate genome-wide profiles of 6mA.^8–10^ Despite its capacity to characterize 6mA in mammalian genomic DNA (gDNA) and mitochondrial DNA (mtDNA), concerns have been raised that the antibody off-target binding, sequence misalignment, and RNA contamination may have led to the artifactual false positive detection of 6mA using these antibody-based methods.^11^ Restriction enzyme digestion-based approaches, such as 6mA-RE-seq and DpnⅠ-seq, yield single-nucleotide resolution modification profiles, but are limited to the detection of specific sequence motifs.^12, 13^

One recent study reported a base-resolution 6mA sequencing method relying on the nitrite-mediated deamination of unmodified A sites.^14^ While the strategy is novel, the method still needs considerable improvement to increase the deamination rate in order to avoid false negative or false positive sites. In addition to the second-generation sequencing-based techniques, the third-generation SMRT sequencing which allows direct recognition of DNA methylation has achieved high-confidence 6mA detection in bacterial genomes.^15^ However, high false discovery rates were observed when performing SMRT on mammalian DNA, possibly due to many factors including the very low levels of 6mA coupled with high 5mC levels, a lack of consistent sequence context for 6mA, and the presence of 6mA-containing bacterial contaminates.^4, 16, 17^

The different limitations of each method mentioned above contributed to the somewhat controversial results of 6mA in the mammalian genome, leaving its presence and functions under debate. The 6mA to A ratio in mitochondrial DNA from human cell lines, for instance, was quantified by UHPLC-QQQ-MS/MS to ∼15 ppm in one study and 130∼400 ppm in another, most likely due to the purity of isolated mtDNA in samples measured.^2, 9, 10^ However, the same cell line exhibited no 6mA enrichment in reads mapped to mtDNA by 6mASCOPE, a newly developed 6mA deconvolution pipeline based on SMRT.^5^ Recognizing the constraints of the current detecting methods, we sought to develop new base-resolution sequencing methods to overcome existing challenges for parsing mammalian genomic features of 6mA.

Our strategy takes advantage of the fact that the methyl group of 6mA destabilizes the Watson–Crick base-pairing although 6mA pairs with deoxythymidine (dT) just like the unmodified deoxyadenosine (dA) .^18, 19^ We reasoned that the use of a thymidine dTTP analog that forms unstable Watson–Crick base-pairing with dA may lead to further weakened base-pairing with 6mA, generating misincorporation signatures opposite of 6mA sites but not A upon DNA replication using selected DNA polymerases, thus allowing base-resolution detection of 6mA. Here, we report the development and application of DR-6mA-seq, which enables high-confidence, mutation-based, base-resolution detection of 6mA in the whole genome. Our results uncovered genomic patterns of 6mA in diverse types of mammalian mtDNA and gDNA. Consistent with previous reports, we observed very low 6mA levels in most mammalian gDNAs. However, we did detect notably elevated gDNA 6mA methylation in specific cell types.

## Results

### Strategy and development of DR-6mA-seq

As a model system to develop DR-6mA-seq, we used a synthetic biotinylated 100-mer DNA oligo harboring an NN-6mA-NN modification site (N=A/G/T/C). To probe whether 6mA can be read as misincorporation signatures under specific primer extension conditions, we tested all the commercially available non-high-fidelity DNA polymerases with different buffer systems. By separating the newly synthesized DNA strand for sequencing, we were able to calculate the 6mA-to-T/G/C mutation ratios at the 6mA sites. *Bst* 2.0 DNA and EpiMark Hot Start Taq DNA Polymerase exhibited observable mutation ratios in some motif contexts, consistent with recent reports that *N*^6^-methylated adenosine could be read as mutations.^20^ We further applied the buffer system of EpiMark Hot Start Taq DNA Polymerase to *Bst* 2.0 DNA and observed an improved mutation proficiency; however, the mutation rates are still too low to be useful.

We reasoned that if we could amplify the base pair stability difference between 6mA versus dTTP and dA versus dTTP, we could increase the mutation frequency opposite of 6mA. We hypothesized that chemically modified dTTPs may enlarge the difference between their base-pairing strength with 6mA and with unmethylated dA. We, therefore, screened all the commercially available dTTP analogs and tested them in the primer-extension system replacing unmodified dTTP. We identified 2-thiothymidine triphosphate (2-thio-dTTP), which carries a sulfur substitution at the 2-position of thymine, which dramatically promoted the misincorporation rate at the 6mA site but not at unmethylated dA sites. This observation is consistent with the diminished base-pairing strength between 6mA versus 2-thio-dTTP and dA versus 2-thiol-dTTP: i) thione does not serve effectively as a hydrogen bond acceptor in Watson–Crick base pairing;^21^ ii) the methyl group on 6mA further weakens the base pairing with canonical 2-thio-dTTP during extension, enabling dATP/dCTP/dGTP to compete with 2-thio-dTTP for base pairing with 6mA (Figure 1A) and causing increased mutation rate at 6mA sites. Meanwhile, dA sites can still base pair with 2-thio-dTTP when using *Bst 2.0* polymerase, generating much fewer mutations (Figure 1A).^22^

**Figure 1.**
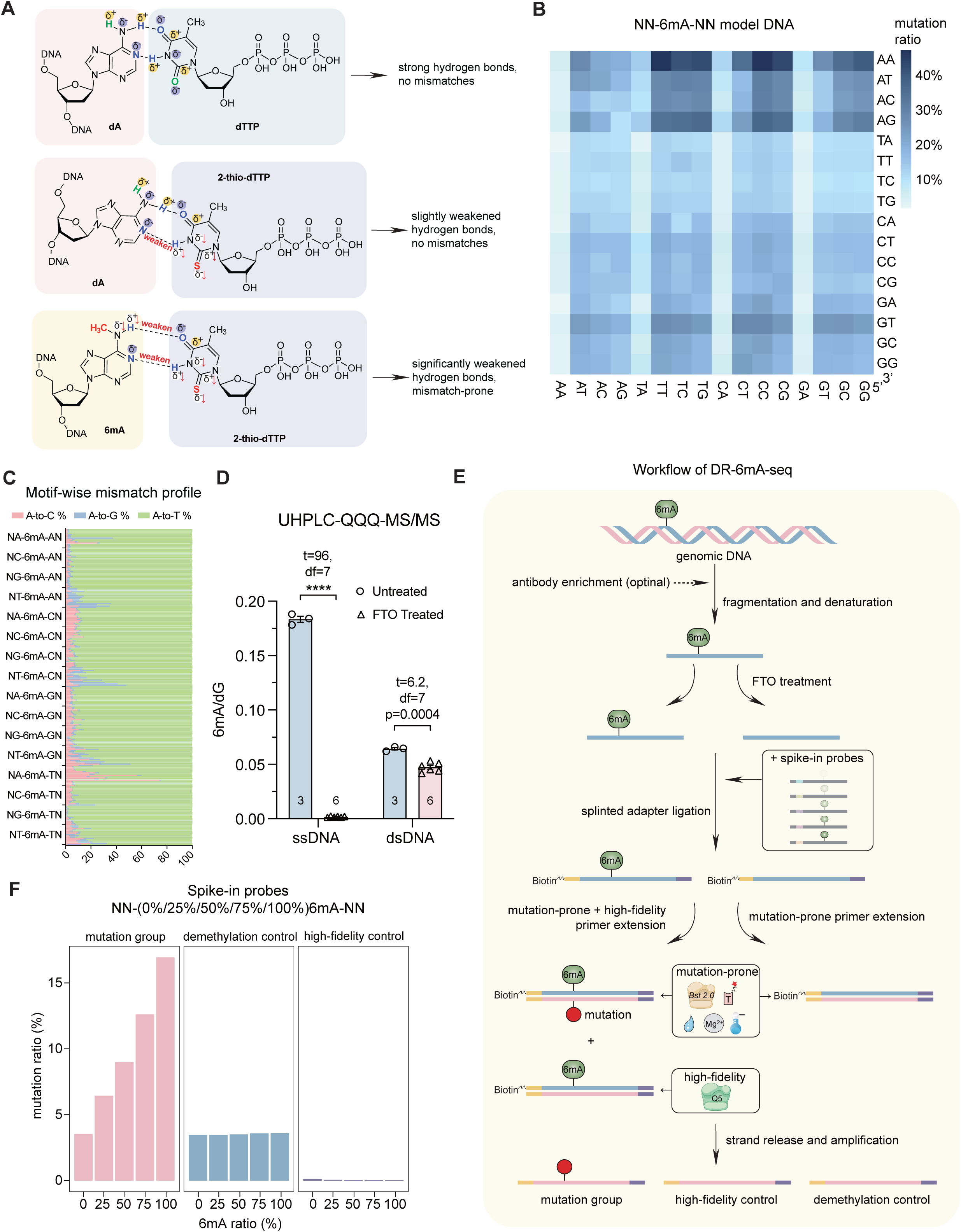
Development and validation of DR-6mA-seq. A) The 6mA-2-thio-dTTP base pair forms weaker hydrogen bonds compared to dA-dTTP and dA-2-thio-dTTP base pairs. B) Heatmap plot for the 6mA-to-T/C/G mutation ratios at 256 motifs (NN-6mA-NN) on the single-stranded model DNA after mutation-prone primer extension. C) The mutation patterns (6mA to T, C, and G respectively) at 256 motifs (NN-6mA-NN) on the single-stranded model DNA. D) FTO demethylates over 99% methylated base of NN-6mA-NN model ssDNA *in vitro* and shows weaker activity on dsDNA. Data are mean ± s.e.m.; analyzed by two-tailed unpaired t-tests. The number of independently repeated reactions are shown in each plot. **** *P* <0.0001. E) Schematic diagram of DR-6mA-seq. Genomic DNA is sequentially denatured, mixed with spike-in probes, treated with FTO (or not), ligated to biotin-tagged adaptors, and extended by *Bst 2.0* DNA polymerase or Q5 high-fidelity DNA polymerase. The synthesized DNA strand is enriched and amplified for next-generation sequencing. 6mA-to-T/C/G mutations were counted for defining 6mA sites and calculating 6mA methylation stoichiometry, using spike-in calibration curves. F) Mutation profiles (including 6mA to T, C, and G) of mutation-prone primer extension on the 6mA spike-in probes harboring 0%, 25%, 50%, 75% and 100% 6mA. Primer extension on FTO-treated probes and primer extension by Q5 high-fidelity DNA polymerase serve as two control groups and do not respond to 6mA fractions. See also **Table S1**.

Having identified the thymidine analog that notably elevates misincorporation opposite of 6mA sites, we further optimized the primer-extension conditions to maximize the mutation ratio. It has long been recognized that the supplement of excess magnesium ions and molecular crowders like gelatins can increase PCR mismatches.^23^ Biased dNTPs ratios and unconventional incubation temperature also facilitate mutations.^24^ We tested a diverse combination of different magnesium ion concentrations, the addition of molecular crowder, dNTP ratios, polymerase concentrations, incubation temperatures, and finally obtained an optimal condition for generating mutation signatures at 6mA sites. Under this condition, the average mutation ratio across 256 NN-6mA-NN motifs is 16%, with the highest mutation ratio at a specific motif reaching 50% (Figure 1B, Table S1). Most of the mutations were 6mA-to-T conversions, the occurrence of which were followed by 6mA-to-C and 6mA-to-G conversions (Figure 1C). These levels of moderate to high mutation rates enable sensitive detection of 6mA in DNA.

### FTO to erase 6mA on ssDNA as a control

To employ this strategy for dissecting genomic features of 6mA, any false positives or mutations induced by other forms of potential modifications must be eliminated. Fat mass and obesity-associated protein (FTO) is known as an efficient eraser of *N*^6^-methyladenosine (m^6^A) and catalyze mRNA m^6^A demethylation in an iron(II)- and α-ketoglutarate-dependent manner.^25^ Previous reports have also shown its biochemical demethylation activity towards 6mA on ssDNA, which could serve as a key control for 6mA identification in DR-6mA-seq.^26, 27^ We purified the recombinant human FTO protein and treated the aforementioned NN-6mA-NN model ssDNA probes, as well as its double-stranded form by annealing with the unmodified complementary strain, with the FTO protein in the presence of iron(II) and α-ketoglutarate (α-KG). After nucleotide digestion and UHPLC-QQQ-MS/MS, we observed that over 99% 6mA on the ssDNA probes was removed by the FTO treatment, while 26.4% of the modifications on dsDNA probes were removed (Figure 1D). We suspect some of the 6mA-containing regions on dsDNA may partially unwind to single-stranded conformation during FTO treatment, which may have led to 6mA demethylation activity observed on dsDNA. Nevertheless, we have confirmed FTO as a powerful ssDNA demethylase of DNA 6mA *in vitro* without significant motif bias. Notably, the *in vitro* efficiency of ALKBH1 as an ssDNA demethylase was reported to be ∼50%.^28, 29^ Thus, the FTO treatment can serve as a critical control for specifically identifying and mapping 6mA in DNA.

Taken together, we formulated a workflow of DR-6mA-seq with one FTO-demethylation control and one high-fidelity control (Figure 1E). RNA-free gDNA or mtDNA was shorn to short single-stranded fragments, and half of the DNA was separated, denatured, and demethylated using FTO. Half of the untreated DNA as well as the demethylated DNA was ligated and then treated with *Bst 2.0* for primer extension, respectively, while the other half of the untreated DNA was extended by using Q5 high-fidelity DNA polymerase as another control. The newly synthesized strands were enriched and amplified for building sequencing libraries. By comparing the mutation profiles of *Bst* 2.0-extended DNA from methylated DNA and FTO-demethylated DNA, and eliminating the background using the high-fidelity control, confident 6mA candidate sites can be uncovered.

To ensure an accurate estimation of the 6mA modification level based on the 6mA-to-T/G/C mutation ratio, we synthesized five spike-in calibration probes that contain known fractions of 6mA (0%, 25%, 50%, 75%, and 100%) at the NN-6mA-NN modification site (Supplementary Table 1-2). Applying DR-6mA-seq to these probes, we observed significant linear correlations (R^2^ > 0.98, p < 0.001) between average mutation frequencies and 6mA fractions, providing a reliable approach to quantify 6mA modification levels within different sequence contexts. Notably, the average mutation rate on unmodified spike-in probes was around 3.5%, which is nearly identical to that of the fully modified spike-in probes after FTO treatment (Figure 1F), confirming that FTO treatment erases 6mA. These calibration probes with unique barcodes were added to each real biological DNA sample when performing DA-6mA-seq procedures for constructing sample-specific calibration curves to minimize any batch effects. The modification fractions for each high-confident 6mA site were estimated by the corresponding linear regression model according to the motif context around the 6mA sites.

### Quantitative 6mA maps of *E. coli* gDNA using DR-6mA-seq

To validate the performance of DR-6mA-seq, we first investigated gDNA from *Escherichia coli* K-12, a wild-type strain with the well-characterized 6mA methylome.^30^ To reveal highly confident 6mA sites, we included 3 biological replicates for DR-6mA-seq (Figure 2A). A total of 20,090 6mA sites were identified (Figure 2B, Table S2), with methylation levels highly reproducible (Figure 2C) in different biological replicates. Most of the detected sites showed ∼100% 6mA modification level, which agrees with the previous reports (Figure 2D).^31^ Notably, 86.7% of the 6mA sites uncovered by DR-6mA-seq overlapped very well with those detected using SMRT sequencing, indicating the high accuracy of our method (Figure 2E).^32^ Further analysis showed 85.2% of the detected 6mA sites fall on the G(6mA)TC motif (Figure 2F), which is the motif recognized by the Dam methylase.^6, 30, 33^ Strikingly, the second most dominant motif of 6mA (1.5%) turned out to be GC(6mA)CNNNNNNGTT, which is the consensus sequence recognized by hsdM, the methylase subunit of EcoKI (Figure 2F).^34, 35^ Our results and the SMRT results agree well on genome-wide 6mA detection, although some of the 6mA sites display lower modification levels using DR-6mA-seq than SMRT (Figure 2G). This is possibly because our approach is more direct and we sampled through more DNA molecules than SMRT.

**Figure 2.**
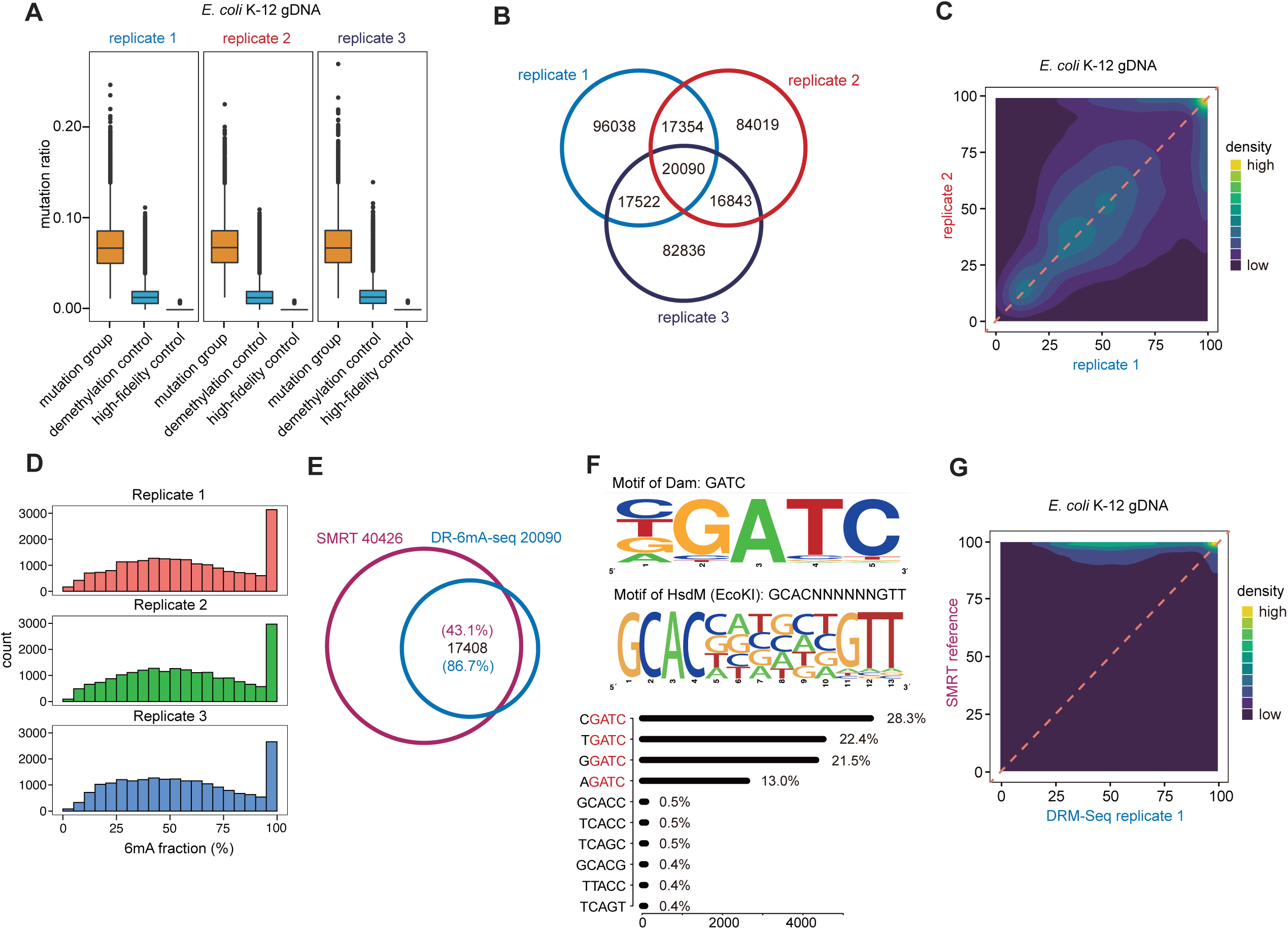
DR-6mA-seq uncovers the quantitative base-resolution 6mA map in *E. coli* genome. A) The box plot of mutation ratio distribution at 6mA sites detected in *E. coli* gDNA, revealed by DR-6mA-seq. The mutation ratios of three groups are shown, i.e., *Bst 2.0* extended DNA from untreated DNA (mutation group), *Bst 2.0* extended DNA from FTO-treated DNA (demethylation control), and Q5-extended DNA from untreated DNA (high-fidelity control). B) Venn diagram showing the overlapped 6mA sites among three biologically independent replicates, detected by DR-6mA-seq. C) Correlation analysis of methylation fractions in *E. coli* K-12 genome, indicating a high correlation between replicates of DR-6mA-seq. D) Histogram indicating the distribution of 6mA fractions at the modified sites in *E. coli* K-12 genome, revealed by DR-6mA-seq. E) Venn diagram showing the excellent overlap between DR-6mA-seq-detected 6mA sites and SMRT-detected 6mA sites in *E. coli* K-12 genome. F) Motif sequence logo and the list of consensus motifs containing 6mA sites in gDNA from *E. coli* K-12, uncovered by DR-6mA-seq. The frequency percentages of the top 5-base 6mA-containing motifs are shown. G) Correlation analysis of 6mA methylation fractions in *E. coli* K-12 genome, indicating a high correlation between 6mA sites detected by DR-6mA-seq and SMRT. See also Tables S2.

### 6mA exists in mouse and human mtDNA

Vertebrate genomes contain much less or base-line level 6mA than prokaryotic ones.^3^ Eukaryotic mitochondrion is known to originate from α-proteobacteria within the bacterial phylum^36^, and we have previously reported the presence of 6mA on mtDNA from human cells.^10^ Our and other studies have shown that 6mA level in HepG2 mtDNA can be 1,300-fold higher than that in gDNA and we have identified 23 6mA sites by ChIP-exo approach.^9, 10^ However, these observations were not supported by the recent SMRT-based 6mASCOPE work and antibody-dependent MM-Seq work, both of which tested mtDNA from HEK293T cells. ^5, 37^ To address this, we performed UHPLC-QQQ-MS/MS using purified HEK293T mtDNA with gDNA digested by exonuclease V. Contrary to the high 6mA abundance in HepG2 mtDNA (400 ppm)^10^, we did not detect 6mA signal in HEK293T mtDNA, which explain the previously reported observation by 6mASCOPE or MM-seq (Figure S1E-I).^5, 6^ We performed DR-6mA-seq with RNA-depleted mtDNA purified from HepG2 cells, as well as adult mouse brain and liver tissues (Figure S1A, Table S3-S5). After context-specific calibration and reproducible filtering in two biological replicates, we identified 159, 58, and 40 overlapped 6mA sites in mtDNA from HepG2 cells, mouse brain, and mouse liver, respectively (Figure 3A). The identified 6mA sites in both species spread across the mitochondrial genome, except for a region in mouse mtDNA that is not uniquely mappable (Figure S1C, S1D).^38^ Different from *E. coli* results, most 6mA sites identified in human and mouse mtDNA showed low modification levels (Figure 3B, Figure S1B), which might explain the weak mtDNA 6mA signals detected by the SMRT-based method. Notably, DR-6mA-seq showed high reproducibility even for some lowly methylated 6mA sites (Pearson r = 0.74 for HepG2 cell line, Pearson r = 0.67 for mouse brain tissue, and Pearson r = 0.78 for mouse liver tissue; Figure 3B).

**Figure 3.**
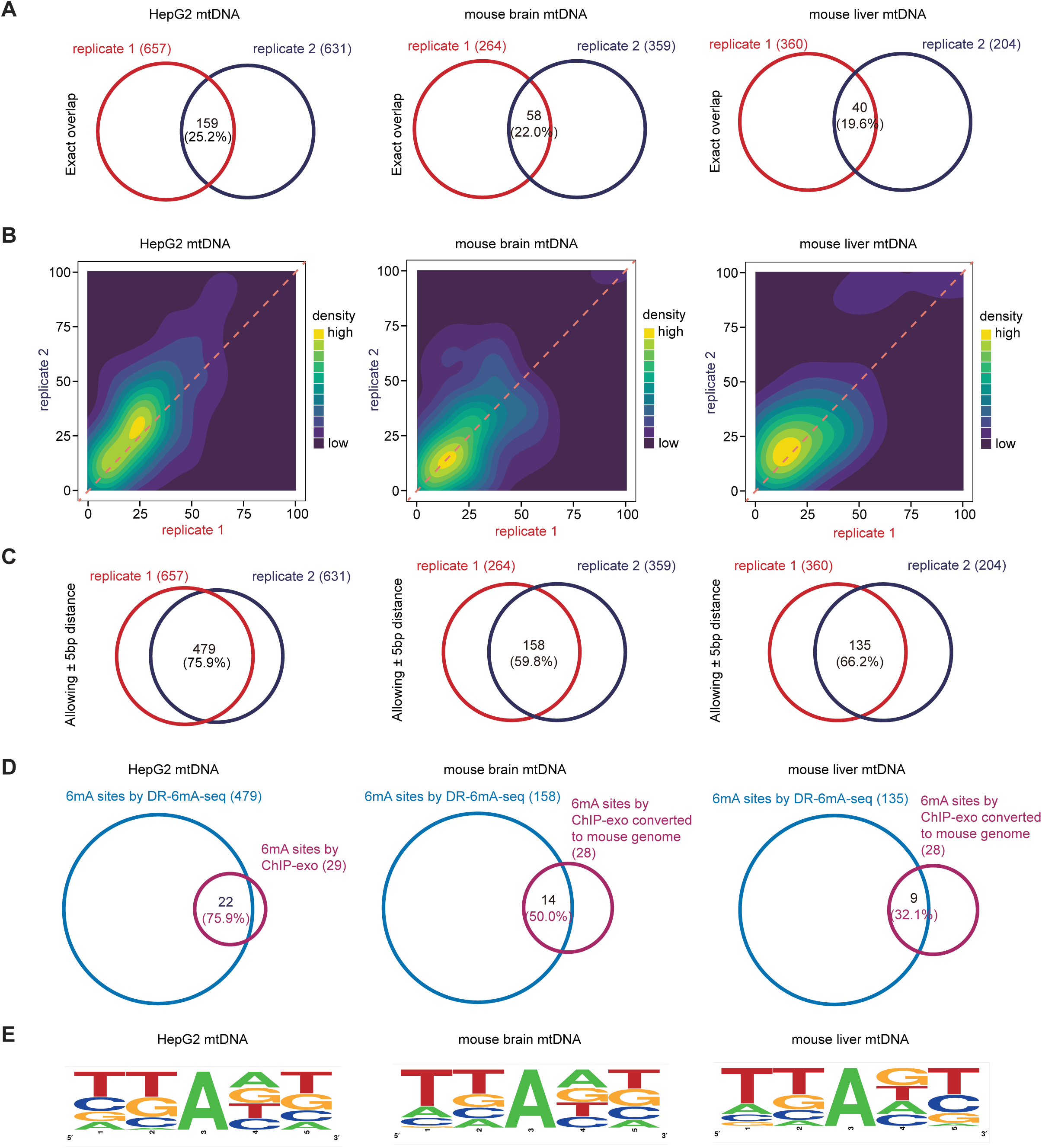
DR-6mA-seq detects widely distributed 6mA in human and mouse mitochondrial DNA. A) Venn diagram showing the overlapped mtDNA 6mA sites detected by DR-6mA-seq between two biologically independent replicates from HepG2 cells, mouse brain, and mouse liver. B) Correlation analysis of 6mA methylation fractions in mtDNA from HepG2 cells, mouse brain and mouse liver, indicating a high correlation between replicates of DR-6mA-seq. C) Venn diagram showing the overlapped mtDNA 6mA sites (allowing ± 5 bp window) between two biologically independent replicates from HepG2 cells, mouse brain, and mouse liver, detected by DR-6mA-seq. D) Venn diagram showing the excellent overlap between DR-6mA-seq-detected 6mA sites and ChIP-exo-detected 6mA sites in mtDNA from HepG2 cells, mouse brain, and mouse liver. E) Motif sequence logo of the consensus motifs containing 6mA sites in mtDNA from HepG2 cells, mouse brain, and mouse liver, uncovered by DR-6mA-seq. See also Figure S1 and Tables S3-S5.

We reasoned that the low methylation levels of mtDNA 6mA sites may suggest a region-based methylation model resembling promoter DNA 5-methylcytosine methylation, instead of strict site-specific methylation; or 6mA methylation may occur to specific regions of mtDNA instead of at specific sites. To test this, we extended all 6mA sites to a window flanking 5 bp to generate lower-resolution maps of mtDNA 6mA sites, and 59.8-75.9% 6mA sites could be reproduced in the two biological replicates (Figure 3C). Further comparison of DR-6mA-seq 6mA sites with our previous ChIP-exo-based 6mA sites also showed high consistency, with 75.9% ChIP-exo-based 6mA sites overlapping with sites uncovered by DR-6mA-seq. Notably, this overlapping percentage is drastically higher than the 5.2% predicted by random overlapping, confirming the existence of 6mA in mtDNA (Figure 3D). Notably, the motif contexts of 6mA sites in mtDNA from different sources showed similar patterns, suggesting evolutionary conservation between mouse and human (Figure 3E).

In summary, we characterized hundreds of 6mA sites in mtDNA using DR-6mA-seq and confirmed an excellent overlap with results from the IP-based method. We showed that 6mA is widely distributed in HepG2 mitochondrial genes with a frequency of over 0.5%, although most of mtDNA 6mA sites display lower modification fractions than those in *E. coli* DNA. The prevalence of 6mA sites detected in mammalian mtDNA supports previously proposed functional roles in mitochondrial transcription and replication.^10^

### 6mA genomic distribution in mouse testis

While emerging evidence including our study above convincingly demonstrates that 6mA is present and plays role in mammalian mtDNA, the existence of 6mA in the mammalian genome has been under debate.^4, 5, 11, 28, 39–45^ To investigate 6mA in the mammalian gDNA, we performed UHPLC-QQQ-MS/MS on gDNA isolated from selected human and mouse tissues and cells. We detected considerable 6mA signals (4∼7 ppm) in the testis gDNA from both human and mouse (Figure 4A, Figure S2A-E). Notably, we did not observe detectable 6mA in the human liver, human brain, or mouse E15.5 embryo gDNA under the same mass spectrometry conditions (Figure S3E-G). In light of this, we analyzed gDNA extracted from 4-month-old mouse testes. Although the LC-MS/MS signal of 6mA from mouse testis gDNA is definitive the level is still quite low. We first applied anti-6mA antibody to enrich 6mA-containing DNA fragments and then performed DR-6mA-seq on the enriched samples (Figure S2F). Sequencing results showed high reproducibility in gDNA replicates (Pearson r = 0.71, Figure 4B), and revealed around 800 methylated sites in each replicate (Figure 4C, Table S6). Among the two biological replicates, although only 106 6mA sites overlapped at the same position, 715 6mA sites can be identified overlapping within the 1-kb flanking windows in both replicates (Figure 4C), supporting region-specific 6mA deposition. These overlapping 6mA sites exist in several main motifs (Figure 4D). While 6mA distributes across the genome, it also shows enrichment at the satellite, rRNA, and tRNA regions (Figure 4F, 4G). Interestingly, a majority of 6mA sites were detected on the Y chromosome in the mouse testis gDNA (Figure 4E). We currently do not know the potential functional relevance.

**Figure 4.**
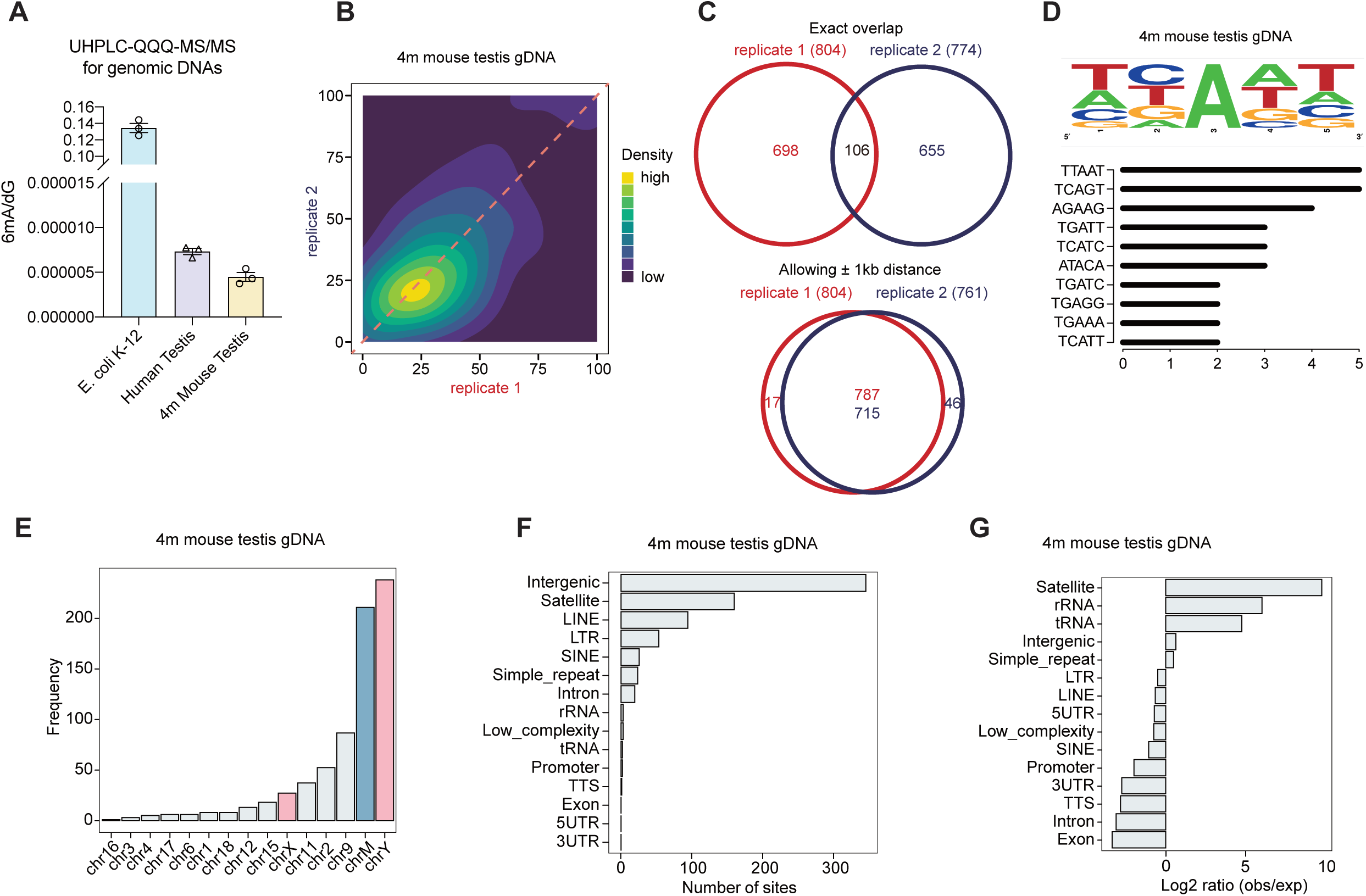
DR-6mA-seq characterizes 6mA modification in the genomic DNA of mouse testis. A) Quantification of 6mA level in gDNA from human and mouse intact testis by LC-MS/MS, with *E. coli* gDNA as a positive control. Data are mean ± s.e.m.. B) Correlation analysis of 6mA methylation fractions in antibody-enriched mouse testis gDNA, indicating a high correlation between replicates of DR-6mA-seq. C) Venn diagram showing the overlapped 6mA sites in mouse testis gDNA, allowing and not allowing ± 1 kb window, detected by DR-6mA-seq between two biologically independent replicates. D) Motif sequence logo and the list of the consensus motifs containing 6mA sites in gDNA from mouse testis, uncovered by DR-6mA-seq. E) Chromosome-wide distributions of gDNA 6mA sites in mouse testis, revealed by DR-6mA-seq. F) Genomic distributions of gDNA 6mA sites in mouse testis, revealed by DR-6mA-seq. G) Genomic enrichment of gDNA 6mA sites in mouse testis, revealed by DR-6mA-seq. See also Figure S2, S3, and Tables S6.

### 6mA is upregulated in *Erbb2*/*neu*\*-transformed NIH/3T3 cells

While we showed 6mA can be present in gDNA in certain mammalian tissues we sought a cell line system that we can perform a more thorough characterization. A recent report using mass spectrometry and antibody-dependent enrichment and sequencing revealed that gDNA 6mA levels could be relatively high in glioblastoma cancer stem cells (GCSs) and primary glioblastoma patient tumors.^28^ We thus selected this system for the further practice of DR-6mA-seq. Among the glioblastoma models, we specifically chose B104-1-1, a frequently used glioblastoma cell line established by transforming NIH/3T3 embryonic fibroblast cells with the activated *Erbb2*/*neu* oncogene (Erbb2 with p.Val661Glu).^46–51^ Taking advantage of their nearly identical genetic background, we decided to investigate potential 6mA changes during gliomagenesis using these two cell lines. We first conducted UHPLC-QQQ-MS/MS with genomic DNA isolated from bacteria- and mycoplasma-free NIH/3T3 and B104-1-1 cell cultures to test their overall gDNA 6mA levels (Figure 5A, S3A-D). The glioblastoma cell line, B104-1-1, harbored a ∼4-fold higher 6mA level (∼80 ppm) than its parental cell line, NIH/3T3. We then enriched 6mA-containing DNA fragments from both B104-1-1 and its parental cell line NIH/3T3 using the anti-6mA antibody and performed DR-6mA-seq.

**Figure 5.**
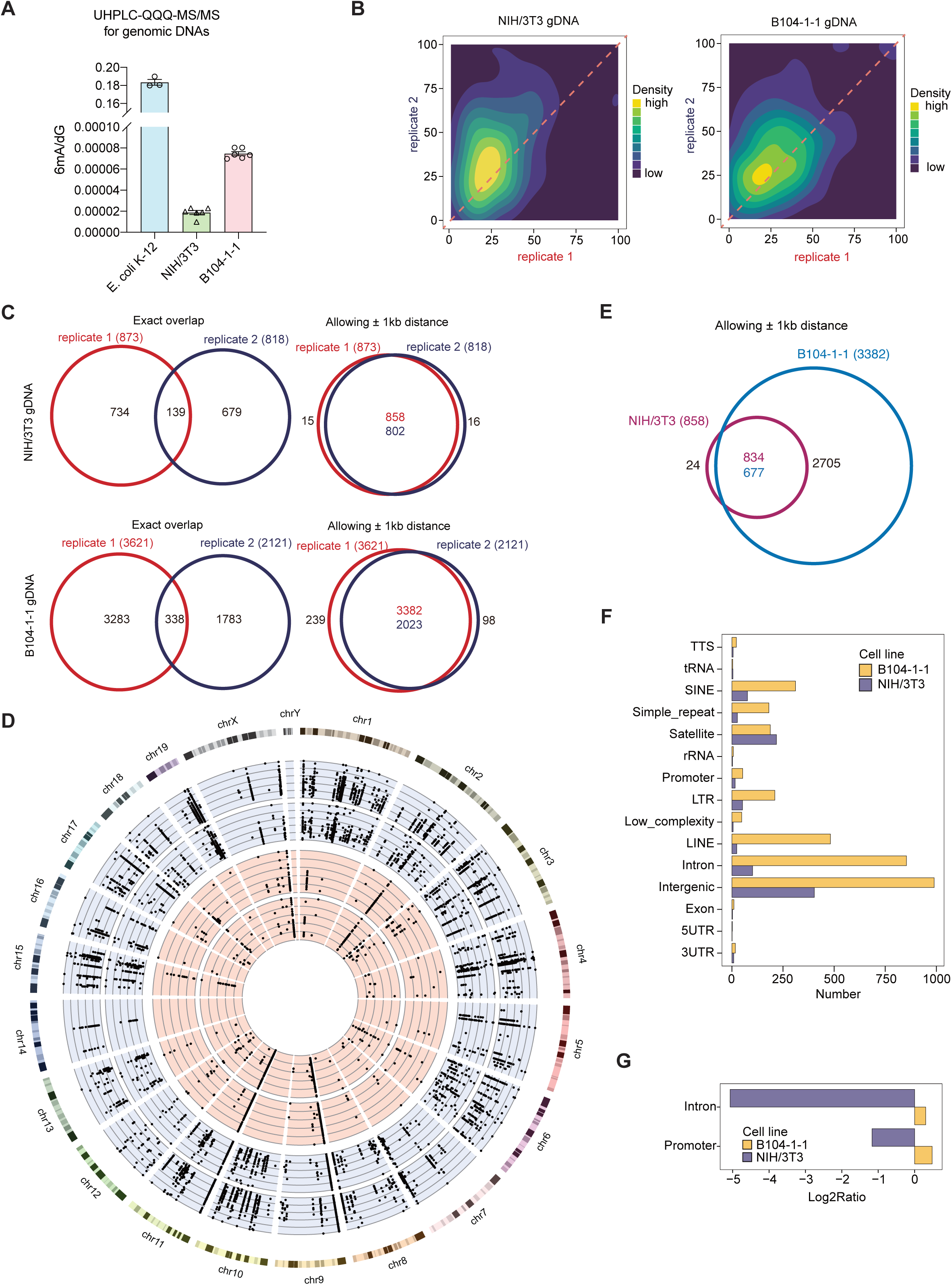
The prevalence of 6mA modification is significantly elevated during the transition from mouse embryonic cells to glioblastoma cells. A) Quantification of 6mA level in gDNA from cultured NIH/3T3 and B104-1-1 cells, by LC-MS/MS, with E. coli gDNA as a positive control. Data are mean ± s.e.m. B) Correlation analysis of 6mA methylation fractions in antibody-enriched gDNA from NIH/3T3 and B104-1-1 cells, indicating a high correlation between replicates of DR-6mA-seq. C) Venn diagram showing the overlapped gDNA 6mA sites in NIH/3T3 and B104-1-1, allowing and not allowing ± 1 kb window, detected by DR-6mA-seq between two biologically independent replicates. D) Distributions of 6mA sites along the mouse genome. The dots on the blue and red background are gDNA 6mA sites in two biologically independent replicates of B104-1-1 and NIH/3T3 cells, respectively, revealed by DR-6mA-seq, with six circled axes representing the methylation fractions at 0%, 20%, 40%, 60%, 80%, and 100%. E) Venn diagram showing the overlapped gDNA 6mA sites between NIH/3T3 and B104-1-1 cells, allowing ± 1 kb window. F) Genomic distributions of gDNA 6mA sites in NIH/3T3 and B104-1-1 cells, revealed by DR-6mA-seq. G) Genomic enrichment of gDNA 6mA sites identified in NIH/3T3 and B104-1-1 cells, revealed by DR-6mA-seq. Enrichment scores on intron and promoter are shown. See also Figure S3-S5 and Tables S7-S8.

We first determined the sex of both cell lines prior to the data analysis (Figure S4A).^52^ Biological replicates gave consistent 6mA sites across the genome, with most 6mA sites exhibiting low or moderate modification fractions (Figure 5B, Table S7-S8). While DR-6mA-seq detected around ∼800-850 gDNA 6mA sites in NIH/3T3 cells, ∼2,000-3,500 6mA sites were observed in B104-1-1 gDNA (Figure 5C, S4B). Approximately 3-fold more 6mA sites were identified in B104-1-1 cells, which is consistent with our mass spectrometry results (Figure 5A). B104-1-1 cells displayed much more unique gDNA 6mA sites on chromosome 1, 10, 19, 4, 15, etc (Figure S4D, S4E). For the two biological replicates, although only 139 and 338 methylated sites overlap exactly at the 6mA site in NIH/3T3 and B104-1-1 gDNA, respectively, above 95% of the 6mA sites overlapped very well within the 1-kb flanking windows (Figure 5C). Although gDNA 6mA methylation in non-cancerous NIH/3T3 cells displayed motif distribution similar to mouse testis, B104-1-1 glioblastoma cells showed different features of 6mA-enriched motifs (Figure S4C). Around 97% of 6mA sites in NIH/3T3 gDNA could be detected in the corresponding glioblastoma cells within the 1-kb flanking windows (Figure 5E). Genomic 6mA sites identified in glioblastoma cells accumulate in the regions of intergenic, intron, promoter, and repeats RNA (LINE, SINE, LTR, etc.) (Figure 5F, 5G, S4F, S4G). Functional enrichment analysis of B104-1-1-specific 6mA sites showed high enrichment of cell projection morphogenesis-related genes (Figure S4H), perhaps reflecting morphology differences between the two cell lines (Figure S4I). Notably, 6mA sites identified in genomic DNA tend to form clusters, which may suggest synergistic effects (Figure 5D, S5A).

To further understand the features of these 6mA sites, we overlapped the gDNA 6mA sites in NIH/3T3 cells with the published ATAC-seq dataset from the same wile-type NIH/3T3 cells,^53^ by setting a sliding window of 500 bp, 1,000 bp or 2,000 bp centered at the 6mA site, respectively. The analysis showed 60-71% of gDNA 6mA sites can overlap very well with ATAC-seq peaks, suggesting a correlation of 6mA with accessible chromatin regions (Figure S5C).^39^ We then overlapped B104-1-1 gDNA 6mA sites with the ATAC-seq peaks in data from glioblastoma tumor induced by implanted GL261 cells,^54^ and observed 10-16% of gDNA 6mA sites overlapping with ATAC-seq peaks (Figure S5D). Most of these overlapped 6mA sites are within ± 1,000 and 1,500 bp ranges around ATAC-seq peak centers in NIH/3T3 cells and B104-1-1 cells, respectively, exhibiting moderate to high modification levels (Figure S5E, S5F), suggesting an association of 6mA with transcriptional activation.

In sum, the application of our method not only confirmed the previous report of the presence of 6mA in glioblastoma cells but also identified 6mA sites for future mechanistic investigation of the exact roles of genomic 6mA.

### Mammalian 6mA validation

Because of the low levels of 6mA in mammalian gDNA, we performed antibody-based enrichment first and then followed with DR-6mA-seq to reveal the presence and modification sites of 6mA. Unlike the mtDNA case in which we obtained the absolute 6mA fraction at each modified site, the results from gDNA enrichment and sequencing only showed the locations of 6mA sites but not modification fraction at the individual site. To quantify the actual modification fraction at the 6mA sites revealed by DR-6mA-seq, we next performed the amplicon sequencing assay targeting the representative 6mA sites. Different from the standard DR-6mA-seq library construction flow, the amplicon assay needs to consider the extension stop occurring at 6mA sites for calculating the actual mutation ratio at each 6mA site (Figure 6A).

**Figure 6.**
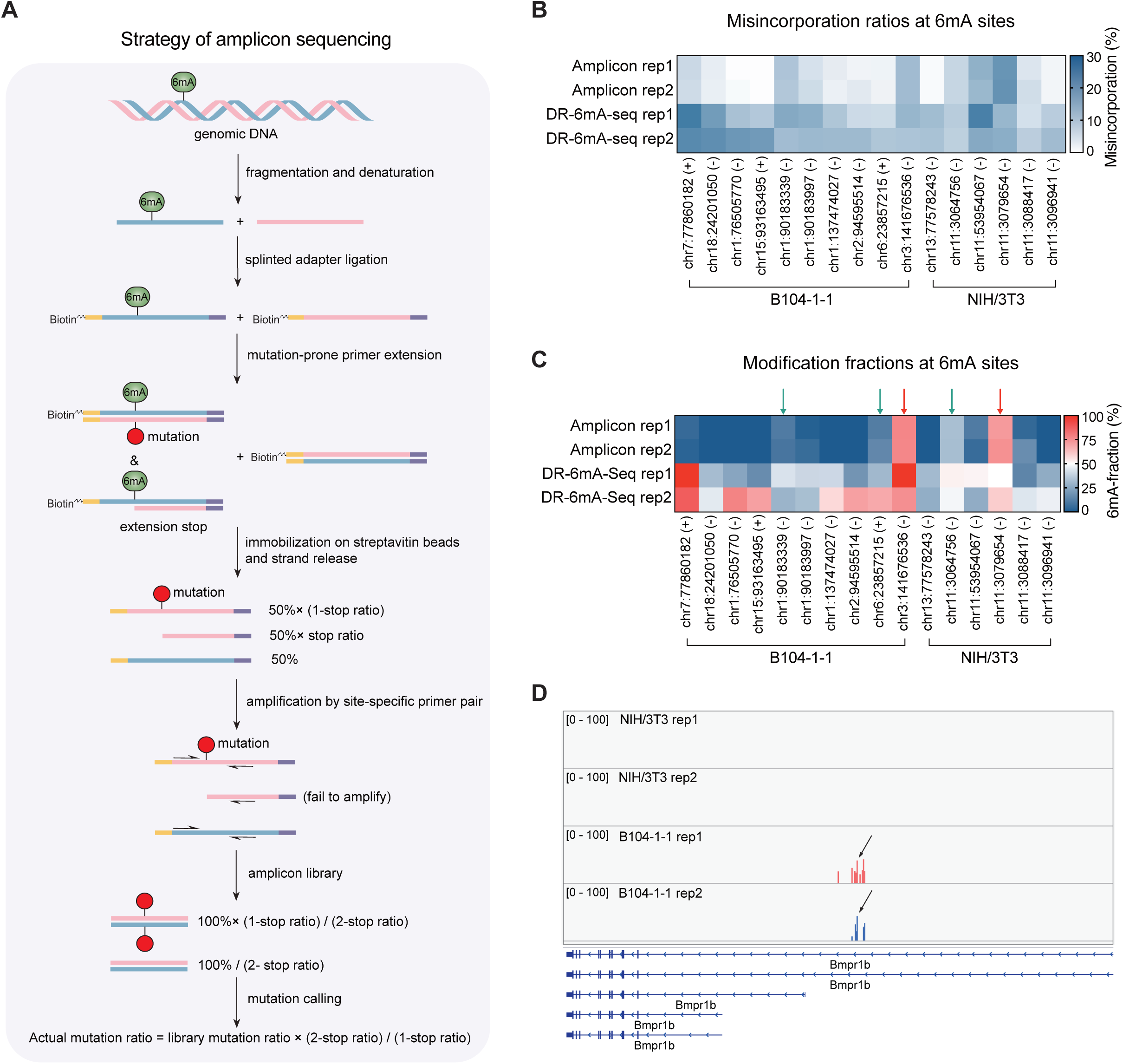
Validation of gDNA 6mA sites by amplicon assay. A) A flowchart of amplicon assay for gDNA 6mA validation, revealing 6mA fraction by adjusted misincorporation ratios. B) The actual misincorporation ratios obtained in amplicon assay versus the misincorporation calculated from DR-6mA-seq data. C) The 6mA methylation fractions by amplicon assay versus 6mA fractions calculated from DR-6mA-seq data. gDNA 6mA sites showing >70% estimated modification in amplicon assay were marked by red arrows; other gDNA 6mA sites showing above 10% modification fraction were labeled by green arrows. For B) and C), 16 gDNA 6mA sites were investigated, with 6 from NIH/3T3 cells and 10 from B104-1-1 cells. D) A representative 6mA cluster located at *Bmpr1b*, with the arrow marking the 6mA site of the highest modification fraction identified in C). See also Figure S5-S6.

From the gDNA 6mA sites detected in NIH/3T3 cells and B104-1-1 cells, we selected 16 6mA sites with surrounding sequences suitable for designing amplicon assay primers (avoiding repeat sequences for instance). We then used gDNA without antibody enrichment for amplicon sequencing to determine the actual 6mA modification fractions. The actual mutation ratios at most 6mA sites were found below 5%, indicating low 6mA fractions. However, for several sites in both cell lines, we observed 5-20% actual mutation ratios in the amplicon assay (Figure 6B). After converting the mutation ratios to 6mA modification fractions using the calibration curves (Figure 1B, 1F), we identified five gDNA 6mA sites displaying above 10% 6mA fraction, with two sites showing ∼70-80% 6mA fractions (Figure 6C). Intriguingly, the highest modified 6mA site (∼87%, chr3:141676536,) we identified, together with the nearby lower-modified sites, falls on *Bmpr1b* in the B104-1-1 genome but not in the NIH/3T3 genome (Figure 6D). We further noticed that *Bmpr1b* is a gene closely associated with the tumorigenicity of glioblastoma and exclusively expressed in B104-1-1 but not in NIH/3T3 (Figure S5B), which may hint at a potential regulatory function of 6mA in the tumorigenicity of B104-1-1.^55, 56^ The amplicon assay provides strong evidence to demonstrate the reliability of DR-6mA-seq and confirmed the existence of 6mA in mammalian gDNA, although only a few sites were shown to accumulate to high fractions that could be potentially functional.

To provide a completely orthogonal method for DNA 6mA validation, we modified a previously reported method based on a silver-ion-mediated base-paring affinity assay.^57^ A proper concentration of Ag^+^ can selectively stabilize the dA–dCTP mismatch and promote primer extension, while the complex of 6mA–Ag^+^–dCTP is unstable and leads to the termination of primer extension. We selected two 6mA sites of the highest modification fraction in HepG2 mtDNA (chrM:5523, tRNA W; chrM:9219, *CO3*), as well as the top 6mA site validated by amplicon assay in the glioblastoma gDNA (chr3:141676536, *Bmpr1b*), and performed the silver-ion-mediated base-paring affinity assay with biotinylated dCTP for both untreated gDNA and FTO-treated gDNA (Figure S6A). Through RT-qPCR quantification, we captured a significantly higher level of biotinylated DNA, which indicates the higher dCTP incorporation ratio after FTO treatment, and thus confirmed the presence of 6mA methylation at these mtDNA and gDNA 6mA sites (Figure S6B).

## Discussion

*N*^6^-methyldeoxyadenosine (6mA) was first discovered in *Bacterium coli* back in the mid-twentieth century.^58^ Over a few decades, 6mA has gradually been recognized as a critical DNA mark in prokaryotes and protists; however, much less is known about its presence and functions in mammals. It is present in the genomes of low eukaryotes and invertebrates, displays distinct distribution patterns, and plays regulatory roles in transcription.^8, 59–63^ Recent studies have also established the presence and potential functional roles of 6mA in mitochondrial DNA in mammals.^9, 10^ In retrospect, it is not a surprise that mtDNA is 6mA modified because of its bacterial origin. However, the presence of 6mA in mammalian genomic DNA has been under intense debate, mostly because of the lack of truly sensitive detection methods. We have proposed early on that 6mA may play limited but regulatory roles in specific cells or during specific biological processes because of its very low levels in mammalian genomes.^64^

Here we present DR-6mA-seq, an antibody-independent, base-resolution sequencing method with high sensitivity to help probe 6mA. Using the FTO-mediated 6mA demethylation control, DR-6mA-seq is also highly specific to 6mA. We have validated the method using *E. coli* gDNA and human mtDNA. We confirmed that the gDNA 6mA levels in most human tissues are too low to be detectable under our LC-MS/MS conditions; however, our mass spectrometry results did confirm visible levels of 6mA in mouse testis gDNA as well as the mouse glioblastoma cell gDNA. By applying DR-6mA-seq, we have the first maps of 6mA distribution in these gDNA. We also applied amplicon sequencing to reveal the exact modification fraction of 6mA sites. As expected, most sites are modified at low levels, but we did observe sites that modified at 70-80% fraction, suggesting potential functional relevance. We also optimized an orthogonal silver-ion-mediated base-paring affinity assay and confirmed results from our DR-6mA-seq.

Our DR-6mA-seq results revealed enrichment of mammalian 6mA in intergenic regions, intronic regions, LINE elements, SINE elements, and LTR (Figure 4F, S4F, S4G), consistent with previous reports using DIP-seq.^41^ Certain 6mA sites are not only conserved across biological replicates but also shared by mouse testis and embryonic fibroblast cells. The overall abundance of mammalian 6mA differs quite dramatically across tissues and can be elevated upon the expression of one single oncogene in the glioblastoma case, with some sites reaching high modification fractions. These results do support the presence of 6mA, albeit limited in modification sites and modification fraction, and potential functional relevance in specific cell types (glioblastoma) or in specific biological processes (development). We believe future applications of DR-6mA-seq can reveal not only 6mA distribution in different genomes of interest but also potential dynamic changes at modification sites to dissect functional implications. Lastly, the use of modified dNTP for elevated misincorporation opposite of a modified base via weakened Watson-Crick base pairing is a strategy that could be applied to detect other nucleic acid base modifications with increased sensitivity.

## Author contributions

C.H. and X.F. conceived the idea and designed the experiments. X.F. and L.-S.Z. performed most of the experiments. X.C. analyzed the high-throughput sequencing data. C.Y., P.W., Y.Z., and Z.Z. provided assistance with the experiments. X.F, X.C., L.-S.Z., and C.H. wrote the manuscript.

## Declaration of interests

C.H. is a scientific founder, a member of the scientific advisory board and equity holder of Aferna Green, Inc. and AccuaDX Inc., and a scientific co-founder and equity holder of Accent Therapeutics, Inc.

## STAR Methods

### RESOURCE AVAILABILITY

#### Lead contact

Further information and requests for resources should be directed to and will be fulfilled by the lead contact, Chuan He.

#### Materials availability

All unique reagents generated in this study are available from the lead contact with a completed Materials Transfer Agreement.

#### Data availability

The raw and processed DR-6mA-seq data have been deposited into NCBI Gene Expression Omnibus (GEO) database with accession number ‘GSE213876’.

### EXPERIMENTAL MODEL AND SUBJECT DETAILS

HepG2, NIH/3T3, and B104-1-1 cells were purchased from ATCC, cultured in DMEM (Gibco, 11965092) with 10% FBS (Gibco, 26140079), and grown at 37°C with 5% CO2. NIH/3T3 from ECACC were maintained in the same condition but only used in sex determination. As a part of cell authentication, cell lines used in this study were examined by mycoplasma contamination test using LookOut Mycoplasma PCR Kit (Sigma, MP0035).

### METHOD DETAILS

#### Development of DR-6mA-seq protocol using synthetic DNA

The 100-mer biotin-tagged NN-6mA-NN model DNA oligo and primer-extension primer (Table S9) was synthesized by IDT (Integrated DNA Technologies). At first, the primer-extension primer was denatured at 70℃ for 2 min and immediately chilled on ice. In a total volume of 50 μL, 50 ng model DNA and 2.5 μL 2 μM oligo-primer-extension primer were mixed with different DNA polymerases, such as *OneTaq* DNA Polymerase, Vent DNA Polymerase, phi29 DNA Polymerase, Pwo SuperYield DNA Polymerase, EpiMark Hot Start *Taq* DNA Polymerase, *Bst* DNA Polymerase-full length, *Bst* 2.0 DNA Polymerase, *Bst* 2.0 WarmStart DNA Polymerase, *Bst* 3.0 DNA Polymerase, Recombinant HIV reverse transcriptase), and the supplied buffers, with the supplement of different concentration of magnesium sulfate, manganese chloride, polyethylene glycol (PEG8000, PEG4000, PEG1000, PEG600, PEG600, PEG200), dATP/dCTP/dGTP and modified dTTP/UTP (dUTP, UTP, 2-Thio-dTTP, 4-Thio-dTTP, 5-Aminoallyl -dUTP, Fluorescein-12-dUTP). Primer extension tests were further carried out under different temperatures. The newly synthesized strands by extension were subsequentially released from the biotinylated template strands immobilized on Streptavidin C1 beads (Thermo Fisher, 65001) by incubation in 150 mM NaOH. The released strands were neutralized, cleaned up by Oligo Clean & Concentrator Kit (Zymo Research, D4061), and incubated with Streptavidin C1 beads again for the complete removal of template strands. The supernatant was cleaned up with Oligo Clean & Concentrator Kit and amplified for 10 cycles by NEBNext Ultra II Q5 Master Mix (NEB, M0544S) and NEBNext Multiplex Oligos for Illumina (NEB, E7335S). The resulting libraries were purified by gel electrophoresis and recovery (NEB, T1020S). All libraries were sequenced at 2M reads depth in single-ended 100 bp mode at NovaSeq 6000 sequencer (Illumina). The primer extension condition was finalized as: 1500 units of *Bst 2.0* DNA polymerase (NEB, M0537M), 1× EpiMark Hot Start Taq Reaction Buffer (NEB, M0490S), a mixture of dATP (NEB, N0446S), dCTP (NEB, N0446S), dGTP (NEB, N0446S), 2-Thio-dTTP (Trilink, N-2035), and 8% PEG4000 (Rigaku, # 25322-68-3). All reactions were incubated at 60℃ for 1 h.

#### Expression and purification of recombinant human FTO protein

As described previously, the human FTO gene (GenBank Accession No. NP_001073901.1) was subcloned into pET28a vector (Novagen) and used for transformation with the BL21 (DE3) *Escherichia coli* strain.^65^ When the optical density at 600 nm reached 1.0, cells were cooled to 16 °C, for 1L bacteria culture, 1 mL 100 mM IPTG, and 200 μL 4 μg/μL (NH₄)₂Fe(SO₄)₂ solution was added, and cells were cultured at 16 °C for an additional 16 h. The cells from 1L culturing media were pelleted and resuspended in 40 mL 1X Lysis Buffer (20 mM Tris-HCl 7.5, 300 mM NaCl, 5 mM imidazole (pH 7.5), 1 tablet of Roche cOmplete Protease Inhibitor Cocktail (EDTA-free)) and sonicated. After centrifugation, the supernatant containing the soluble recombinant protein was filtered through 0.22 um syringe filters and then purified with Ni Sepharose 6 Fast Flow (GE Healthcare). After washing the agarose once with 50 mL 1X Lysis Buffer, wash the agarose with 50 mL of two washing buffers respectively (Wash Buffer A: 20 mM Tris-HCl 7.5, 500 mM NaCl, 5 mM imidazole (pH 7.5), 1 tablet of Roche cOmplete Protease Inhibitor Cocktail (EDTA-free); Wash Buffer B: 20 mM Tris-HCl 7.5, 300 mM NaCl, 25 mM imidazole (pH 7.5), 1 tablet of Roche cOmplete Protease Inhibitor Cocktail (EDTA-free)). The protein was eluted in 14 mL Elution Buffer (20 mM Tris-HCl 7.5, 300 mM NaCl, 250 mM imidazole (pH 7.5)). The collected flowthrough was initially concentrated by centrifuge and then diluted by 1X Resuspension Buffer (20 mM Tris-HCl pH 7.5, 300 mM NaCl) and re-concentrated 5 times to remove imidazole. The protein was finally concentrated to ∼22 mg/mL using Ultra 0.5 Centrifugal Filters (Millipore, UFC503096). Glycerol was added in the enzyme to a final concentration of 25% and stored at –80 °C for future use.

#### Preparation of DNA samples

Genomic DNA from *Escherichia coli* strain K-12 was purchased from ATCC (10798D-5). The human brain, liver, heart, and testis gDNA were purchased from Zyagen (HG-201; HG-314; HG-801; HG-401). Mitochondrial DNA of HepG2 cells, mouse brain, and mouse liver was prepared using Mitochondrial DNA Purification Kit (Abcam, ab288088). Genomic DNA of NIH/3T3 cells, B104-1-1 cells, 4-month-old male C57BL/6J mouse whole testis, 4-month-old male C57BL/6J mouse whole brain, and 2.5-month-old female C57BL/6J mouse whole brain were all prepared using Monarch Genomic DNA Purification Kit (NEB, T3010S). All the DNA samples were harshly treated with RNase A (NEB, T3018L) again prior to library construction to get rid of any RNA contaminants.

#### LC-MS/MS analysis

##### Validation of FTO demethylation activity on ssDNA

The 100-mer biotin-tagged NN-6mA-NN model DNA oligo was annealed to its complementary oligo (Table S9) in Nuclease Free Duplex Buffer (IDT, 11-05-01-12) with the mole ratio of 1:3. 300 ng denatured NN-6mA-NN model ssDNA oligo, as well as 300 ng annealed NN-6mA-NN model dsDNA, was incubated at 37 ℃ for 4 hours in 500 μL reaction solution containing 50 mM HEPES pH 7.0, 60 mM KCl, 75 μM ammonium iron(II) sulfate hexahydrate, 2 mM L-asorbic acid, 300 μM α-ketoglutarate and 2.4 μM recombinant FTO protein. The demethylation reactions were then cleaned up by Oligo Clean & Concentrator Kit (Zymo research, D4061) and DNA Clean & Concentrator Kits (Zymo research, D4003) respectively, and then digested with Nucleoside Digestion Mix (NEB, M0649S) at 37 ℃ for 2 hrs. The digested DNA was filtered through 0.22 μm filter (Millipore, SLGVR04NL). For UHPLC–QQQ–MS/MS analysis, 10 μL sample was injected, and the nucleosides were separated by reverse-phase UHPLC on a C18 column (Agilent, 927,700-092) followed by MS detection using Agilent 6460 QQQ–MS/MS set to multiple reaction monitoring (MRM) in positive electrospray ionization mode. Nucleosides were quantified using the nucleoside precursor ion to base ion mass transitions of 266.1-150.0 for 6mA and 268.1-152.0 for dG. The concentration of nucleosides was quantified using the calibration curves which were obtained from nucleoside standards running at the same condition.

##### Quantification of gDNA 6mA in biological samples

Cell cultures have been treated by plasmocin (InvivoGen, ant-mpt) to ensure zero contamination of mycoplasma. Genomic DNA from cell lines (such as NIH-3T3 and B104-1-1 cells) and tissues (such as *E. coli* K-12, mouse testis, mouse E15.5 embryo, human liver, human brain, human testis) were digested by Nucleoside Digestion Mix (NEB, M0649S) with 1X Reaction Buffer (50 mM potassium acetate pH 5.4, 1 mM ZnCl_2_) at 37 ℃ for 2 hrs. For UHPLC–QQQ–MS/MS analysis, 10 μL sample was injected, followed by the aforementioned standard procedures with a revised UHPLC protocol. To enhance 6mA signal at mass spec and achieve a better separation from other peaks, we adjusted the UPHLC protocol as: 98% A + 2% B at 0 min; 97% A + 3% at 8 min; 93% A + 7% B at 9 min; 89% A + 11% B at 10min; 50% A + 50% B at 10.25∼12.50 min; 98% A + 2% B at 12.75 min; A = H_2_O + 0.1% HCOOH, B = MeOH + 0.1% HCOOH.

#### Optimized DR-6mA-seq protocol for biological DNA samples

For *E. coli* DNA and mtDNA, 400 ng∼1 μg RNA-free DNA was diluted by TE buffer (Invitrogen, AM9849) with 0.1 M NaOH and sonicated to 100-300 nt length using Bioruptor Pico sonication device. For mammalian gDNA, antibody enrichment was performed prior to libraries construction: 50∼100 μg RNA-free DNA was diluted by TE buffer (Invitrogen, AM9849) with 0.1 M NaOH and sonicated to 100∼300 nt length. The fragmented DNA was immunoprecipitated using 25 μg anti-6mA antibody (Sigma, ABE572 or Abcam, ab151230) by gently rotating at 4℃ overnight. Pierce Protein A Magnetic Beads (Thermo Scientific, 88845) were washed and used to pulldown antigen-antibody complex. Subsequentially, the beads were treated by proteinase K (NEB, P8111S) and the 6mA-enriched DNA was then purified with DNA Clean & Concentrator Kit-25 (Zymo research, D4033). The fragmented DNA was mixed with 15% (mass percentage) spike-in probes (Table S9) and then split into three in the proportion of 3:4:1, each for mutation group, demethylation control, and high-fidelity control. For the demethylation control, DNA fragments were denatured and then incubated at 37 ℃ for 4 hours in a 300 μL reaction containing 50 mM HEPES 7.0, 60 mM KCl, 75 μM ammonium iron (II) sulfate hexahydrate, 2 mM L-asorbic acid, 300 μM α-ketoglutarate and 2.4 μM FTO purified protein. The demethylation reaction was then treated with proteinase K (NEB, P8111S) and cleaned up by Oligo Clean & Concentrator Kits (Zymo research, D4061). The three groups of DNA were denatured and subjected to splint ligation following a previous protocol (Table S9).^66^ The ligation products of mutation group and demethylation control underwent primer extension at 60℃ for 3 hours in the 50 μL reaction containing 6 μM biosample-primer-extension primer (Table S9), 1500 units of *Bst 2.0* DNA polymerase (NEB, M0537M), 1× EpiMark Hot Start Taq Reaction Buffer (NEB, M0490S), a mixture of dATP (NEB, N0446S), dCTP (NEB, N0446S), dGTP (NEB, N0446S), 2-Thio-dTTP (Trilink, N-2035), and 8% PEG4000 (Rigaku, # 25322-68-3). Meanwhile, the ligation product of high-fidelity control underwent primer extension in the 50 μL reaction containing 6 μM biosample-primer-extension primer and 1×Q5 High-Fidelity 2X Master Mix (NEB, M0492S) with 6 thermal cycles. The synthesized strands for each group were subsequentially released from the biotin-tagged template strands immobilized on Dynabeads MyOne Streptavidin C1 (Invitrogen, 65001) by incubation in 0.15 M sodium hydroxide. The released strands were neutralized, cleaned up with Oligo Clean & Concentrator Kits, and again incubated with Dynabeads MyOne Streptavidin C1 for the complete removal of the template strands. The supernatant was cleaned up with Oligo Clean & Concentrator Kits and amplified for 10∼12 cycles by KAPA HiFi Uracil+ Master Mix (Roche, KK2801) and NEBNext Multiplex Oligos for Illumina (NEB, T1020S). The resulting libraries were purified by gel electrophoresis and recovery and finally sequenced at 60∼100M depth with single-ended 100 bp mode.

#### Genetic sex determination of NIH/3T3 and B104-1-1 cell culture

The method for sex determination of mouse cell lines was modified from the previous study.^67^ Genomic DNA was isolated from C57BL/6 female and male brain, NIH/3T3 cell cultures (ATCC, CRL-1658, and Sigma 93061524-1VL), and B104-1-1 cell culture (CRL-1887) and amplified by Q5 High-Fidelity 2X Master Mix (NEB, M0492S) and *Rbm31x/y*-F/R primers (Table S9).

#### Quantification of 6mA fraction on specific sites by amplicon sequencing

Genomic DNA was purified from NIH/3T3 cells and B104-1-1 cells and treated by revised DR-6mA-seq protocol to generate mutations at 6mA sites during extension. Without any adaptor ligation, the treated gDNA was used or amplicon assay. For 6mA candidate sites of high mutation ratios or high modification fractions in NIH/3T3 and B104-1-1 cell lines, a 500-nt window was set for designing amplicon primers. The amplicon primers covered the region of 150-300 nt, with the target 6mA site inside this region. The Illumina sequencing barcodes were embedded into the amplicon primers. Around 1-2 ng treated gDNA was then used for amplicon PCR amplification for 40 cycles under the calculated best annealing temperature of each primer pair. The PCR reaction mixture was then purified by agarose gel, and size selection was performed to cut the target band representing the PCR product from the specific 6mA site region. All libraries were sequenced at NextSeq 500 as paired-end 300bp mode. Sequencing reads were mapped to the 500-nt region allowing 10 mismatches per read. The stop ratio at the 6mA site was determined by comparing the reads coverage of two strands distinguished by Illumina sequencing. The actual mutation ratio at the 6mA site was comprehensively calculated by the mutation ratio and stop ratio observed in the amplicon assay.

#### Validation of 6mA sites in the mammalian genome by Ag^+^ based method

Prepare ssDNA from HepG2 mtDNA, B104-1-1 total DNA, and NIH/3T3 total DNA, with FTO demethylation treatment versus the control. Mix 10 μL ssDNA (∼ 150 ng/μL) with 1 μL 50 μM extension primer, 1 μL 2 mM Biotin-dCTP (Lumiprobe, 2715), 5 μL 10X Standard Taq Reaction Buffer (NEB, M0273S), 5 μL 10 μL freshly prepared AgNO_3_, 0.25 μL Taq DNA Polymerase (NEB, M0273S) and 27.75 μL nuclease-free water. Mix well and incubate at 95℃ for 30s. Repeat 95℃ for 15s, 62 ℃ for 1 min, 68℃ for 10 min for 20 cycles. The reaction mixture was purified by Oligo Clean & Concentrator (Zymo Research, D4061). Elute DNA with 40 μL nuclease-free water. Add 5 μL 10 nM biotinylated ssDNA spike-in oligo into the eluted DNA. Mix well and save 2 μL as ‘Input’. The rest 43 μL DNA was mixed with 30 μL Streptavidin C1 beads and incubated at 4 ℃ for 30 min. Then remove the supernatant and wash the beads with C1-Wash Buffer (50 mM HEPES pH7.3, 300 mM NaCl, 0.05% NP40) seven times. Proteinase K treatment was conducted to elute ssDNA from the beads. The eluted ssDNA was further purified by Oligo Clean & Concentrator (Zymo Research, D4061), as ‘Pulldown’. Both ‘Input’ and ‘Pulldown’ samples were measured by RT-qPCR assays to quantify the enrichment folds of the target 6mA region, in FTO demethylation versus the control.

### QUANTIFICATION AND STATISTICAL ANALYSIS

#### Data processing for whole-genome DR-6mA-seq

Raw sequencing reads were first trimmed to remove adaptors and low-quality nucleotides by Trim_Galore. High-quality reads were first aligned to spike-in sequences by bowtie2 to assess the library quality and mutation levels in different sequence contexts. Reads in high-quality libraries were then aligned to reference genomes by bowtie2 with uniquely mapped reads retained for the following analyses. Samtools was used to remove PCR duplicates.

For mutation calling of mtDNA, only reads mapped to mtDNA were retained. For mutation calling of genomic DNA, reads aligned to mtDNA were removed for the following analyses.

#### Statistical calling of 6mA and assessing FDR of whole-genome DR-6mA-seq

Mutated A or T sites were identified by VarScan software with default parameters. To avoid potential artifacts caused by low sequencing depth, only mutations with more than 20 x depth in all replicate groups were retained. We defined candidate 6mA sites based on the following criteria: 1) Not mutated in the control group; 2) Baseline mutated in the demethylated group; 3) Mutated in the error-prone mutation group in all replicates. A total of 256 linear regression models were generated for all 256 nucleotide contexts based on mutation ratios of spike-in sequences. The methylation levels were predicted based on certain linear models according to the corresponding sequence context.

Comparison of different replicates was generated by Bedtools. For region-level comparison, 6mA sites were first extended to include flanking 5-bp for mtDNA and 1000-bp for gDNA by Bedtools, and then compared through intersectBed of Bedtools. Circos map was used for track visualization. 6mA clusters were visualized by IGV (version 2.6.3).

#### Statistics

Statistical comparisons between two groups were conducted by the unpaired two-tailed t-tests. Statistical comparisons among multiple groups were conducted by one-way ANOVA tests. All statistical analysis and data graphing were done in Prism (version 8.4.0) software.

### Supplemental item titles

**Table S1**: Mutation ratios on 256 motifs of model ssDNA (-NN6mANN-) after mutation-prone primer extension, related to **Figure 1**.

Table S2: 6mA sites identified in *E. coli* K-12 DNA using DR-6mA-seq, related to Figure 2. Table S3: 6mA sites identified in HepG2 mtDNA using DR-6mA-seq, related to Figure 3. Table S4: 6mA sites identified in mouse brain mtDNA using DR-6mA-seq, related to Figure 3.

**Table S5**: 6mA sites identified in mouse liver mtDNA using DR-6mA-seq, related to **Figure 3**.

**Table S6**: 6mA sites identified in mouse testis gDNA using DR-6mA-seq, related to **Figure 4**. **Table S7**: 6mA sites identified in NIH/3T3 gDNA using DR-6mA-seq, related to **Figure 5**.

**Table S8**: 6mA sites identified in B104-1-1 gDNA using DR-6mA-seq, related to **Figure 5**. **Table S9**: Model DNA, adaptors, and primers used in this study, related to Key Resources Table.

## KEY RESOURCES TABLE

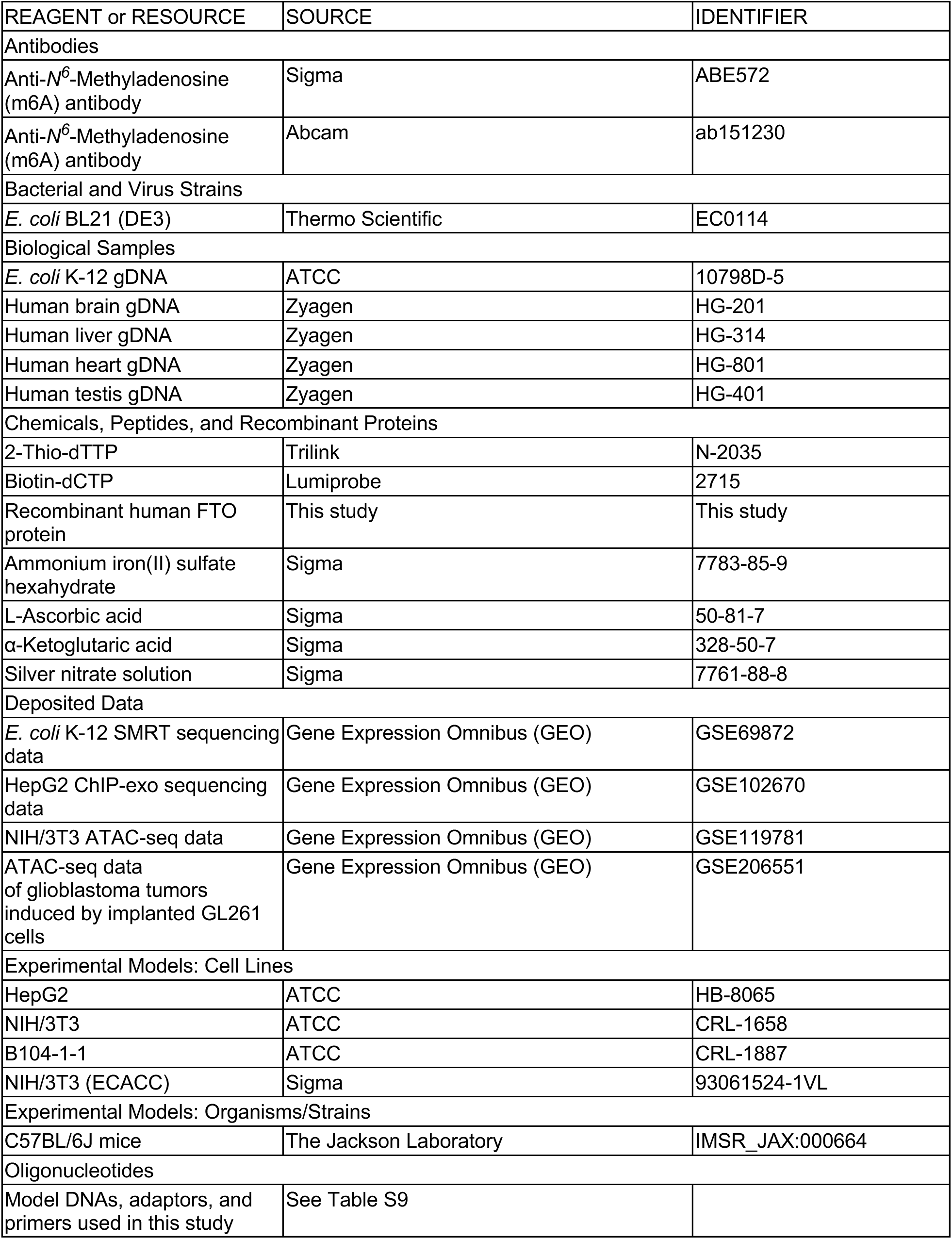
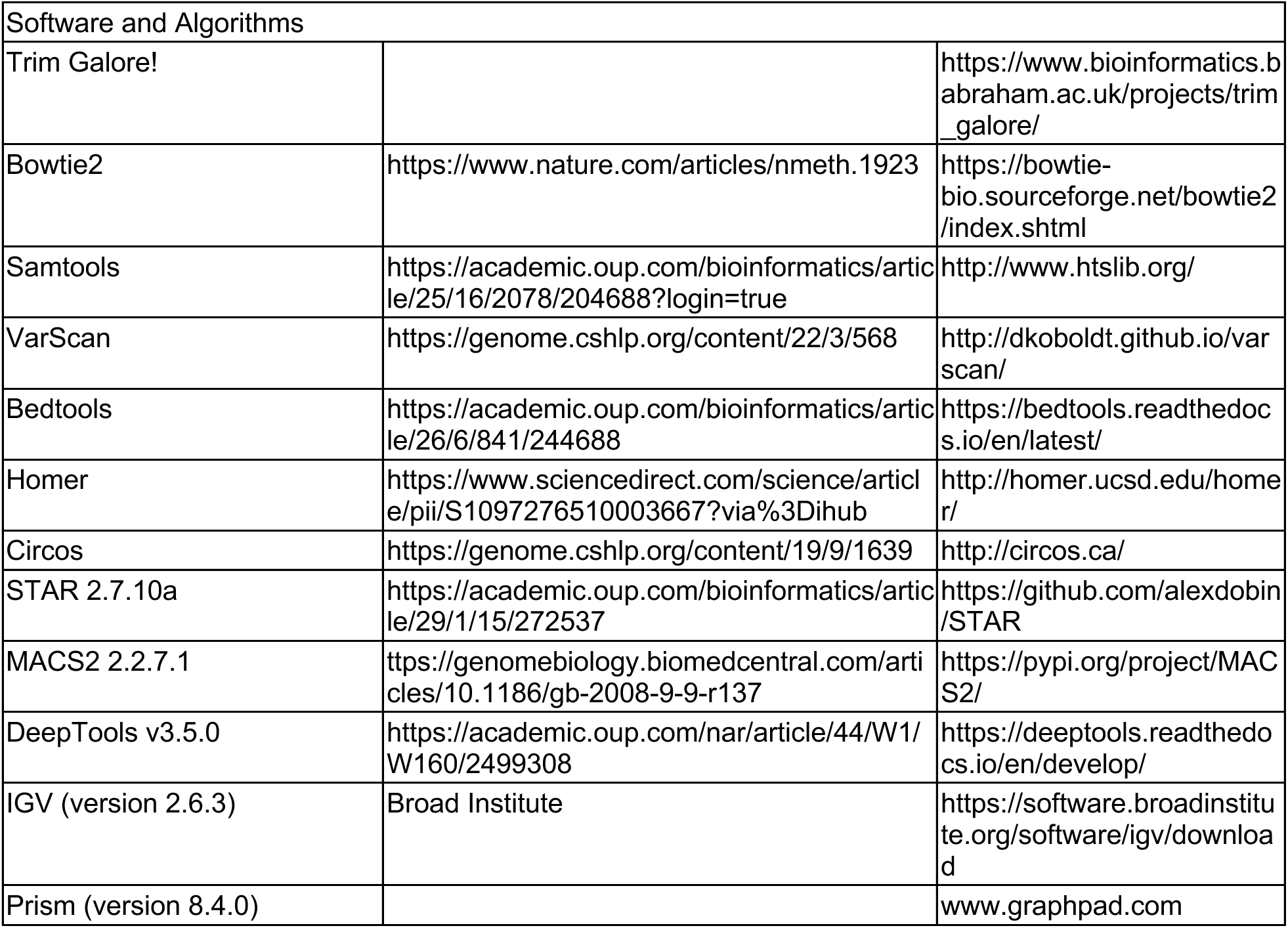

## Acknowledgments

This work was supported by funding from the National Institutes of Health (HG006827 and HG008935) to C.H., and the International Student Fellowship (UChicago, Biological Sciences Division) to X.F.. We appreciate the kind support from UChicago Genomics Facility, Mass Spectrometry Facility, and ARC, especially Dr. C. Jin Qin and Dr. Pieter W. Faber for assisting in the experiments. C.H. is an Investigator of the Howard Hughes Medical Institute.

---

**Figure S1.**
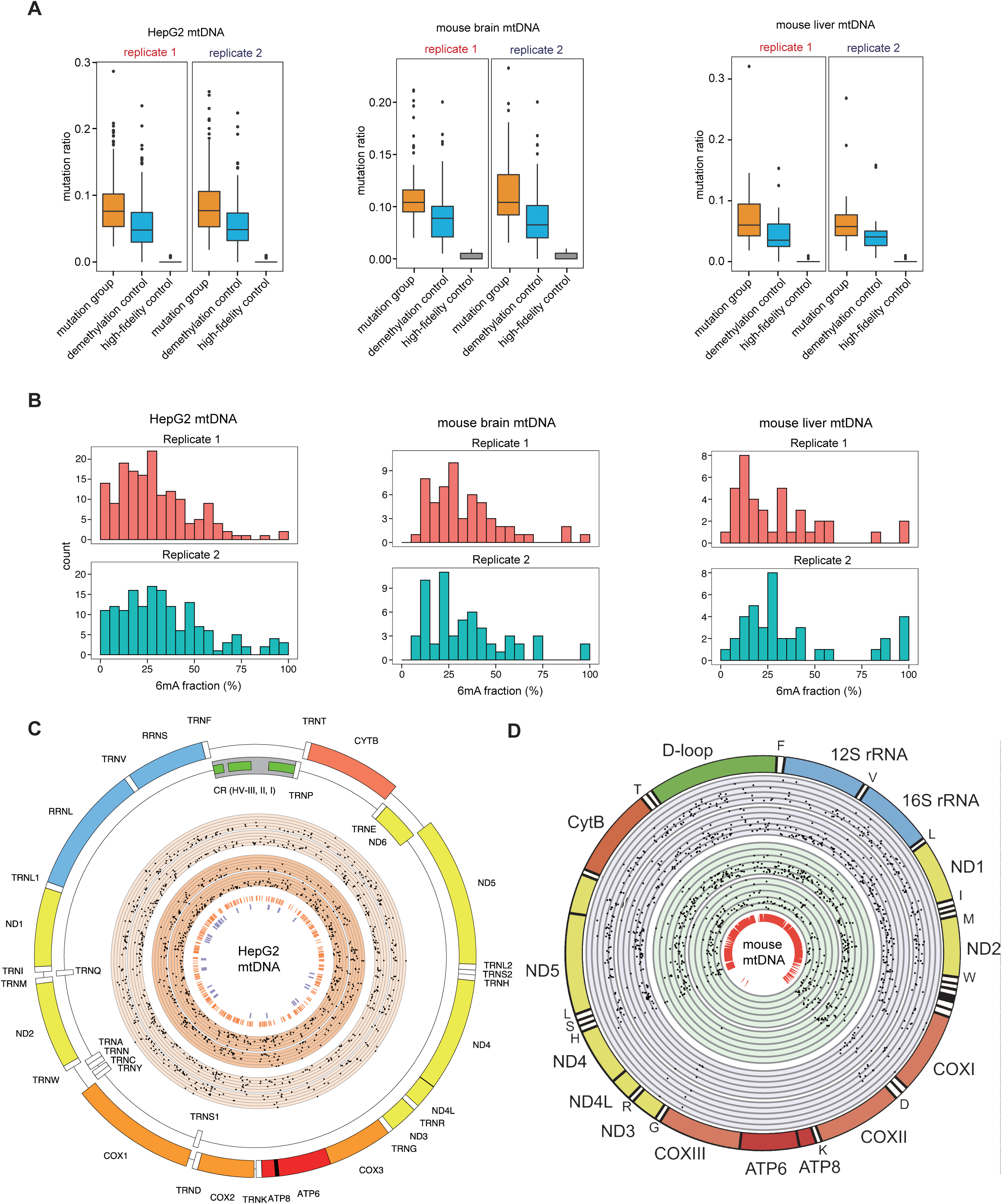
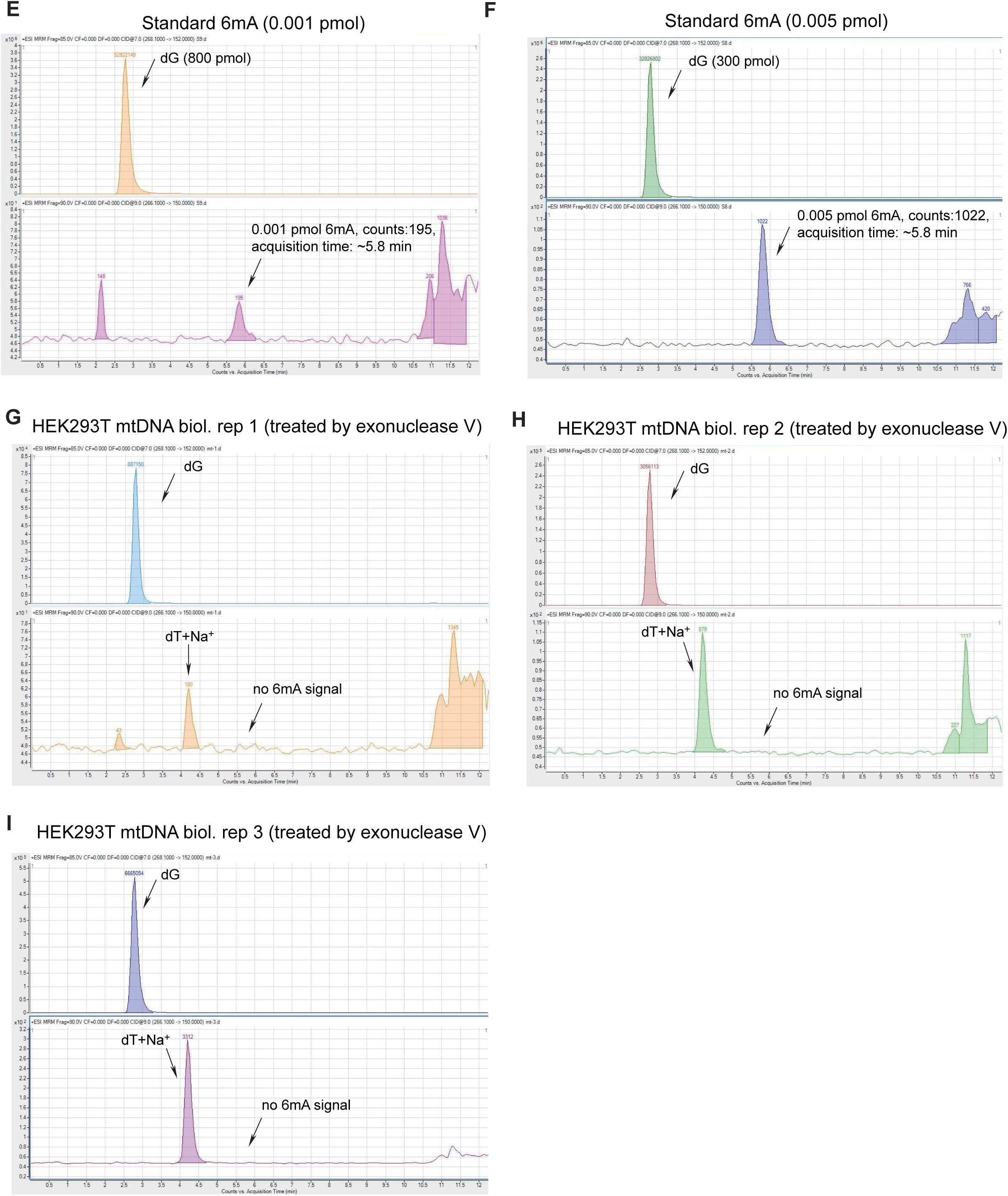
Features of 6mA sites in mtDNA of HepG2, mouse liver, and mouse brain, related to Figure 3. A) The box plot of mutation ratio distribution at 6mA sites detected in HepG2 mtDNA, mouse brain mtDNA, and mouse liver mtDNA, revealed by DR-6mA-seq. The mutation ratios of three groups are shown, such as *Bst 2.0*-extended DNA from untreated DNA (mutation group), *Bst 2.0*-extended DNA from FTO-treated DNA (demethylation control), and Q5-extended DNA from untreated DNA (high-fidelity control). n = 2, biologically independent replicates. B) The histogram of mtDNA 6mA site number distribution normalized to 6mA fraction bins, in HepG2 cells, mouse brain, and mouse liver. n = 2, biologically independent replicates. C) Distribution of 6mA sites along the human HepG2 mitochondrial genome. The dots on the lighter orange background are mtDNA 6mA sites on the forward and reverse strand of mtDNA of HepG2 replicate 1, while the dots on the darker orange background are mtDNA 6mA sites on the forward and reverse strand of mtDNA of HepG2 replicate 2, both revealed by DR-6mA-seq. The six circled axes representing the methylation fractions at 0%, 20%, 40%, 60%, 80%, and 100%. The inner bright orange track represents overlapped mtDNA 6mA sites with 5-bp flanking windows. The inner purple track represents 6mA sites detected by ChIP-exo. D) Distribution of 6mA sites along the mouse mitochondrial genome. The dots on the purple and green background are mtDNA 6mA sites in two biological replicates of mouse brain and mouse liver, respectively, revealed by DR-6mA-seq, with six circled axes representing the methylation fractions at 0%, 20%, 40%, 60%, 80%, and 100%. The inner red track indicates the regions of uniquely mapped reads in mouse mitochondria genome. E) Chromatogram of a representative injection of 6mA (0.001 pmol) and dG standard compound (800 pmol), using LC-MS/MS. The 6mA quantification was calculated by the 6mA peak area, which displays a retention time of ∼5.8 min. F) Chromatogram of a representative injection of 6mA (0.005 pmol) and dG standard compound (300 pmol) using LC-MS/MS. The 6mA quantification was calculated by the 6mA peak area, which displays a retention time of ∼5.8 min. G) Chromatogram of a representative injection of gDNA-depleted HEK293T mtDNA (the first biological replicate), using LC-MS/MS. The optimized LC protocol can distinguish the 6mA peak from the dT+Na^+^ peak which appears at ∼4.2 min. H) Chromatogram of a representative injection of gDNA-depleted HEK293T mtDNA (the second biological replicate), using LC-MS/MS. The optimized LC protocol can distinguish the 6mA peak from the dT+Na^+^ peak which appears at ∼4.2 min. I) Chromatogram of a representative injection of gDNA-depleted HEK293T mtDNA (the third biological replicate), using LC-MS/MS. The optimized LC protocol can distinguish the 6mA peak from the dT+Na^+^ peak which appears at ∼4.2 min.

**Figure S2.**
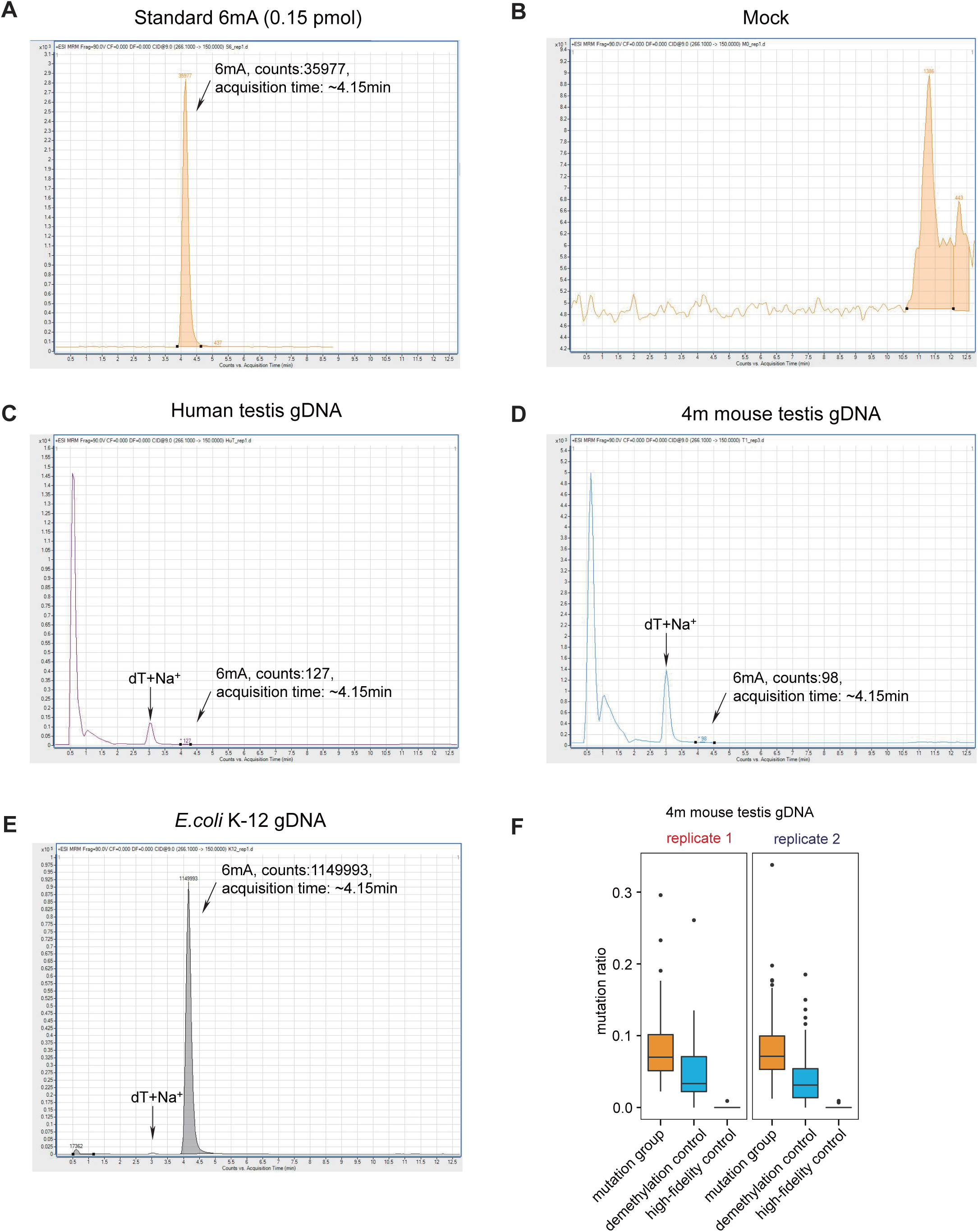
The quantification of gDNA 6mA level by LC-MS/MS and 6mA mutation profiles mapped by DR-6mA-seq in mouse testis, related to Figure 4. A) Chromatogram of a representative injection of 6mA standard compound, using LC-MS/MS. B) Chromatogram of a representative injection of the mock (nucleoside digestion mixture without adding any DNA) using LC-MS/MS. C) Chromatogram of a representative injection of digested human testis gDNA, using LC-MS/MS. The 6mA quantification was calculated by the 6mA peak area, which displays a retention time of ∼4.15 min. The optimized LC protocol distinguished the 6mA peak from the dT+Na^+^ peak which appears at ∼3.00 min. D) Chromatogram of a representative injection of digested mouse testis gDNA, using LC-MS/MS. E) Chromatogram of a representative injection of *E. coli* K-12 gDNA, using LC-MS/MS. F) The box plot of mutation ratio distribution at 6mA sites detected in mouse testis gDNA, revealed by DR-6mA-seq. The mutation ratios of three groups are shown, such as *Bst 2.0*-extended DNA from untreated DNA (mutation group), *Bst 2.0*-extended DNA from FTO-treated DNA (demethylation control), and Q5-extended DNA from untreated DNA (high-fidelity control).

**Figure S3.**
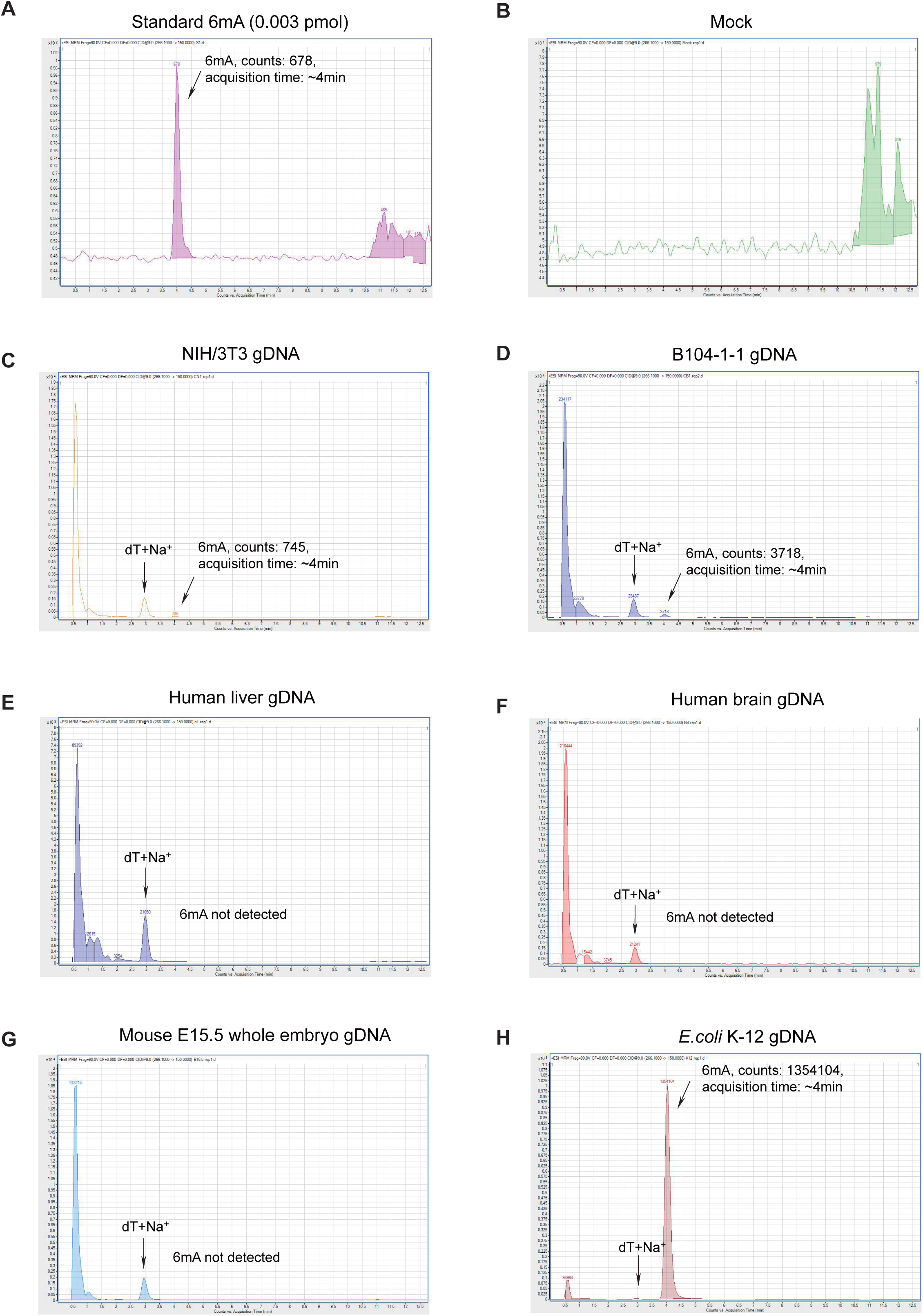
The quantification of 6mA level in gDNA from NIH/3T3, B104-1-1, and multiple mammalian tissues, using LC-MS/MS, related to Figures 4 and 5. A) Chromatogram of a representative injection of 6mA standard compound, using LC-MS/MS. B) Chromatogram of a representative injection of the mock (nucleoside digestion mixture without adding any DNA) using LC-MS/MS. C) Chromatogram of a representative injection of digested gnotobiotic NIH/3T3 (ATCC) gDNA, using LC-MS/MS. The 6mA quantification was calculated by the 6mA peak area, which displays a retention time of ∼4.15 min. The optimized LC protocol distinguished the 6mA peak from the dT+Na^+^ peak which appears at ∼3.00 min. D) Chromatogram of a representative injection of digested gnotobiotic B104-1-1 (ATCC) gDNA, using LC-MS/MS. E) Chromatogram of a representative injection of digested human liver gDNA, using LC-MS/MS. F) Chromatogram of a representative injection of digested human brain gDNA, using LC-MS/MS. G) Chromatogram of a representative injection of digested mouse E15.5 whole embryo gDNA, using LC-MS/MS. H) Chromatogram of a representative injection of *E. coli* K-12 gDNA, using LC-MS/MS.

**Figure S4.**
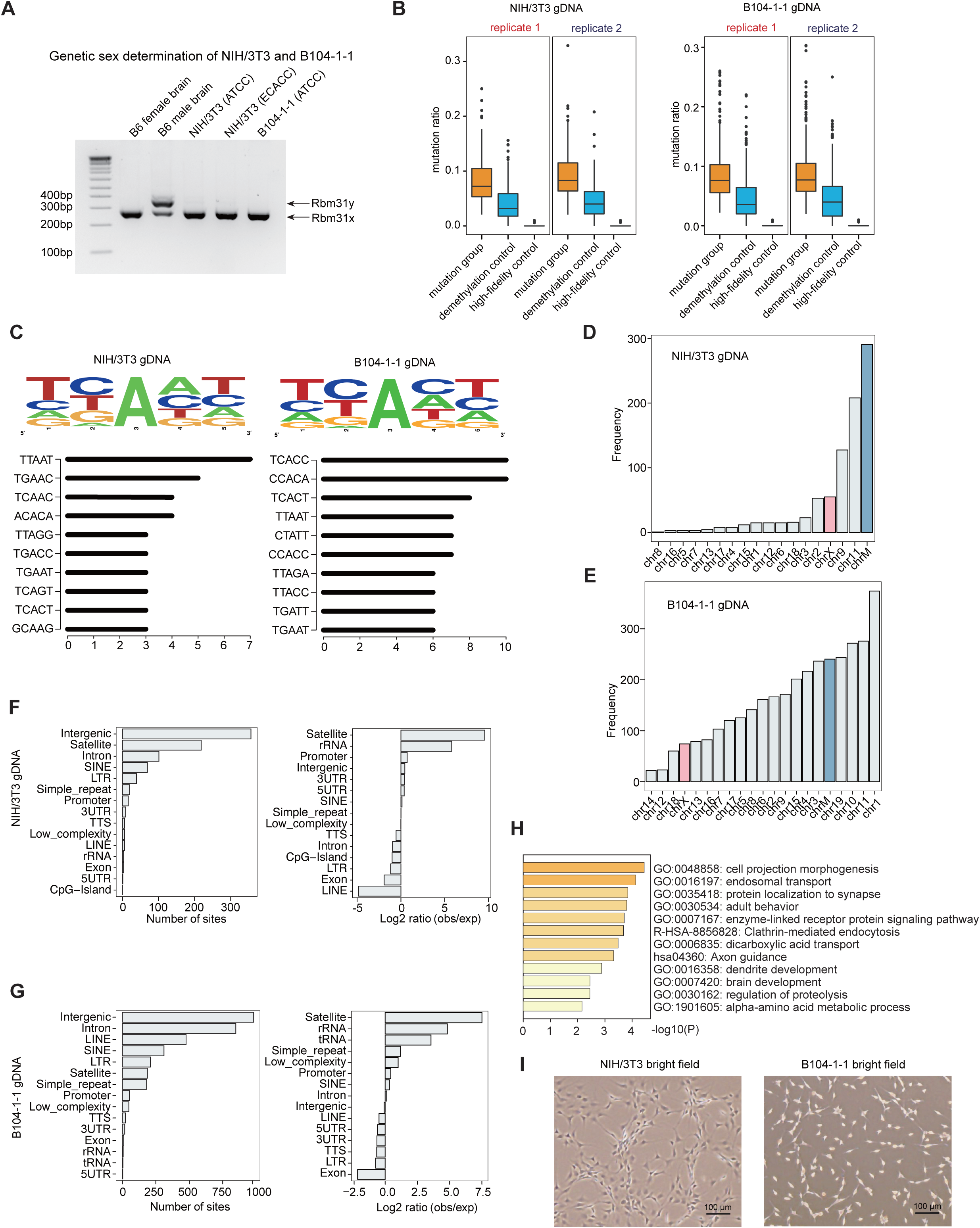
Features of gDNA 6mA sites in NIH/3T3 and B104-1-1 cells, identified by DR-6mA-seq, related to Figure 5. A) Genetic sex determination of NIH/3T3 and B104-1-1 cell lines by amplification of *Rbm31x* and *Rbm31y* by simplex PCR, with female and male mouse brains as controls. B) The box plot of mutation ratio distribution at 6mA sites detected in gDNA from NIH/3T3 (ATCC) and B104-1-1 (ATCC) cells, revealed by DR-6mA-seq. The mutation ratios of three groups are shown, such as *Bst 2.0* extended DNA from untreated DNA (mutation group), *Bst 2.0* extended DNA from FTO-treated DNA (demethylation control), and Q5-extended DNA from untreated DNA (high-fidelity control). C) Motif sequence logo and top 10 consensus motifs containing 6mA sites in gDNA from NIH/3T3 (ATCC) and B104-1-1 cells, uncovered by DR-6mA-seq. D) Chromosome-wide distributions of 6mA sites in gDNA from NIH/3T3. E) Chromosome-wide distributions of 6mA sites in gDNA from B104-1-1. F) Gene annotation shows the genomic distributions and enrichment of gDNA 6mA sites in NIH/3T3 cell lines, revealed by DR-6mA-seq. G) Gene annotation shows the genomic distributions and enrichment of gDNA 6mA sites identified in B104-1-1 cell lines, revealed by DR-6mA-seq. H) The enriched GO clusters of 6mA-modified genes in B104-1-1 cells. Genes with 6mA sites on exons, introns, and promoters are considered as 6mA-modified genes. I) Bright field photographs of NIH/3T3 and B104-1-1 cells.

**Figure S5.**
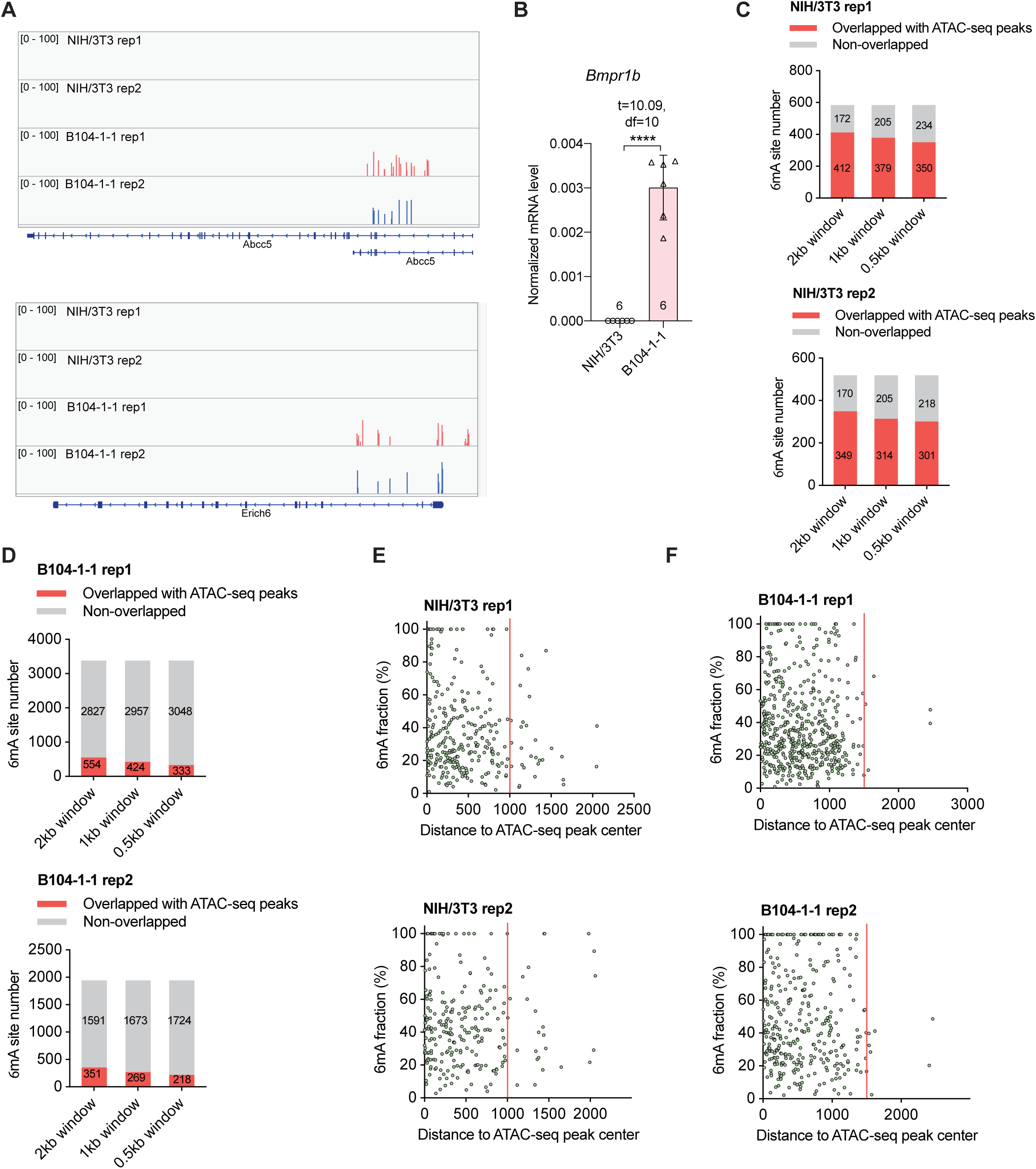
6mA sites identified in NIH/3T3 and B104-1-1 cells tend to form clusters, exhibiting overlaps with ATAC-seq peaks, related to Figure 5. A) Two representative cell-line-specific gDNA 6mA clusters located at *Abcc5* and *Erich6* in B104-1-1 cells. B) RT-qPCR quantification of *Bmpr1b* expression level in NIH/3T3 and B104-1-1 cells, normalized to *Actb*. Data are mean ± s.e.m.; analyzed by two-tailed unpaired t-tests. The number of independently repeated reactions is shown in each plot. **** *P* <0.0001. C) The NIH/3T3 gDNA 6mA site number overlapped with ATAC-seq peaks from wild-type NIH/3T3 cells (GSE119781), by setting a sliding window of 500 bp, 1,000 bp, or 2,000 bp centered at the 6mA site revealed by DR-6mA-seq. D) The B104-1-1 gDNA 6mA site number overlapped with ATAC-seq peaks from glioblastoma tumor induced by implanted GL261 cells (GSE206551), by setting a sliding window of 500 bp, 1,000 bp, or 2,000 bp centered at the 6mA site revealed by DR-6mA-seq. E) The 2-D plot of NIH/3T3 gDNA 6mA methylation fraction versus the distance to ATAC-seq peak center (GSE119781), most 6mA sites distribute within ±1,000 bp range around ATAC-seq peak center. F) The 2-D plot of B104-1-1 gDNA 6mA methylation fraction versus the distance to ATAC-seq peak center (GSE206551), most 6mA sites distribute within ±1,500 bp range around ATAC-seq peak center.

**Figure S6.**
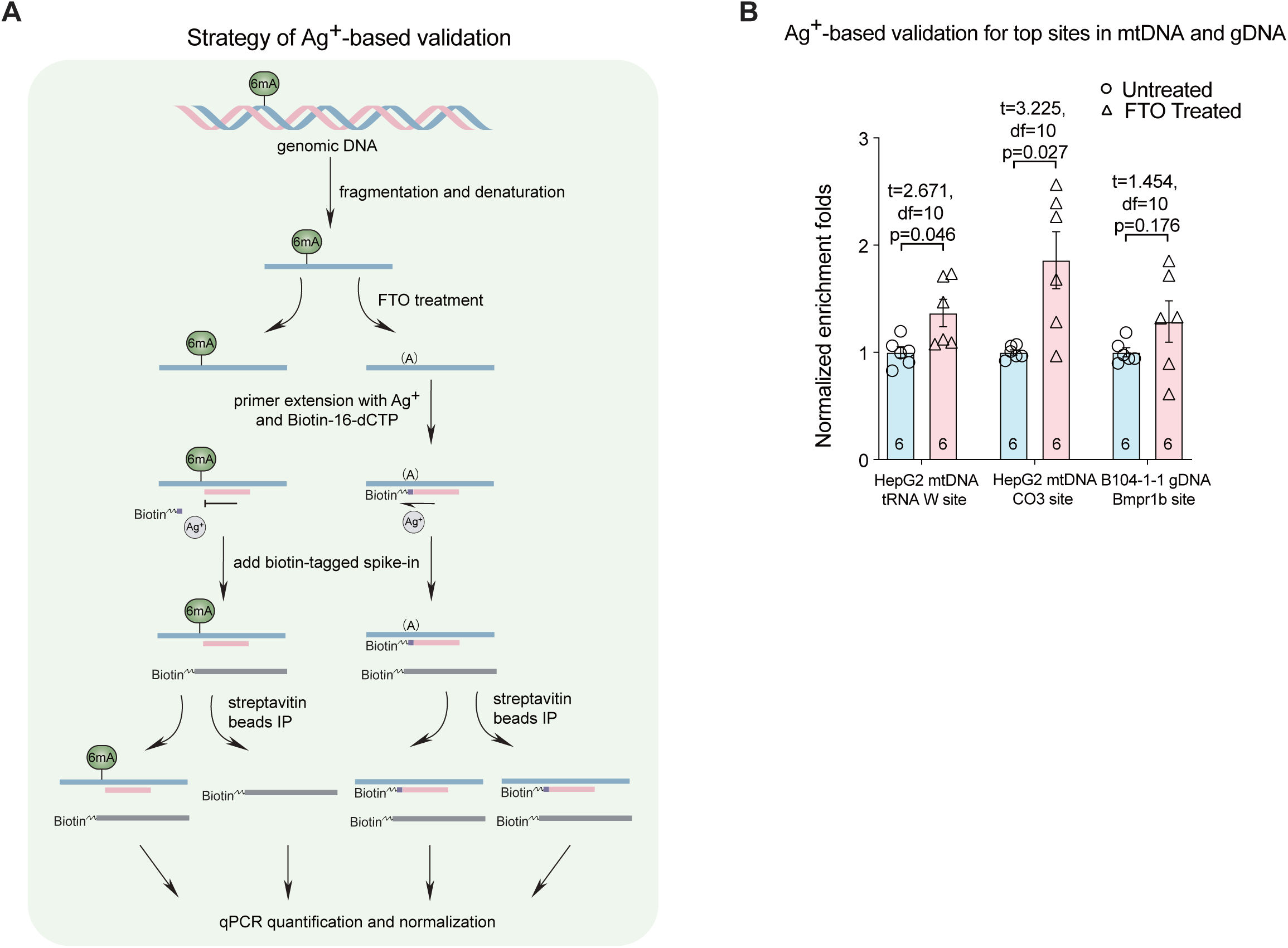
Validation of highly modified 6mA sites in mammalian gDNA by silver-ion-mediated base-paring affinity assay, related to Figure 6. A) Schematic diagram of silver-ion-mediated base-paring affinity assay for 6mA site validation. B) The normalized enrichment folds at highly modified 6mA sites, measured by RT-qPCR-assisted silver-ion-mediated base-paring affinity assay, in the presence and absence of FTO treatment. Data are mean ± s.e.m.; analyzed by two-tailed unpaired t-test. The number of independently repeated reactions is shown in each plot.

## References

1. Liyanage, V., Jarmasz, J., Murugeshan, N., del Bigio, M., Rastegar, M., and Davie, J. (2014). DNA Modifications: Function and Applications in Normal and Disease States. Biology (Basel) 3, 670–723. 10.3390/biology3040670.

2. Shen, C., Wang, K., Deng, X., and Chen, J. (2022). DNA N6-methyldeoxyadenosine in mammals and human disease. Trends in Genetics 38, 454–467. 10.1016/j.tig.2021.12.003.

3. Boulias, K., and Greer, E.L. (2022). Means, mechanisms and consequences of adenine methylation in DNA. Nat Rev Genet 23, 411–428. 10.1038/s41576-022-00456-x.

4. Douvlataniotis, K., Bensberg, M., Lentini, A., Gylemo, B., and Nestor, C.E. (2020). No evidence for DNA N6-methyladenine in mammals. Sci Adv 6. 10.1126/sciadv.aay3335.

5. Kong, Y., Cao, L., Deikus, G., Fan, Y., Mead, E.A., Lai, W., Zhang, Y., Yong, R., Sebra, R., Wang, H., et al. (2022). Critical assessment of DNA adenine methylation in eukaryotes using quantitative deconvolution. Science (1979) 375, 515–522. 10.1126/science.abe7489.

6. O’Brown, Z.K., Boulias, K., Wang, J., Wang, S.Y., O’Brown, N.M., Hao, Z., Shibuya, H., Fady, P.-E., Shi, Y., He, C., et al. (2019). Sources of artifact in measurements of 6mA and 4mC abundance in eukaryotic genomic DNA. BMC Genomics 20, 445. 10.1186/s12864-019-5754-6.

7. Schiffers, S., Ebert, C., Rahimoff, R., Kosmatchev, O., Steinbacher, J., Bohne, A.-V., Spada, F., Michalakis, S., Nickelsen, J., Müller, M., et al. (2017). Quantitative LC-MS Provides No Evidence for m6dA or m4dC in the Genome of Mouse Embryonic Stem Cells and Tissues. Angewandte Chemie International Edition 56, 11268–11271. 10.1002/anie.201700424.

8. Fu, Y., Luo, G.-Z., Chen, K., Deng, X., Yu, M., Han, D., Hao, Z., Liu, J., Lu, X., Doré, L.C., et al. (2015). N6-Methyldeoxyadenosine Marks Active Transcription Start Sites in Chlamydomonas. Cell 161, 879–892. 10.1016/j.cell.2015.04.010.

9. Koh, C.W.Q., Goh, Y.T., Toh, J.D.W., Neo, S.P., Ng, S.B., Gunaratne, J., Gao, Y.-G., Quake, S.R., Burkholder, W.F., and Goh, W.S.S. (2018). Single-nucleotide-resolution sequencing of human N6-methyldeoxyadenosine reveals strand-asymmetric clusters associated with SSBP1 on the mitochondrial genome. Nucleic Acids Res 46, 11659– 11670. 10.1093/nar/gky1104.

10. Hao, Z., Wu, T., Cui, X., Zhu, P., Tan, C., Dou, X., Hsu, K.-W., Lin, Y.-T., Peng, P.-H., Zhang, L.-S., et al. (2020). N6-Deoxyadenosine Methylation in Mammalian Mitochondrial DNA. Mol Cell 78, 382–395.e8. 10.1016/j.molcel.2020.02.018.

11. Lentini, A., Lagerwall, C., Vikingsson, S., Mjoseng, H.K., Douvlataniotis, K., Vogt, H., Green, H., Meehan, R.R., Benson, M., and Nestor, C.E. (2018). A reassessment of DNA-immunoprecipitation-based genomic profiling. Nat Methods 15, 499–504. 10.1038/s41592-018-0038-7.

12. Bird, A.P., and Southern, E.M. (1978). Use of restriction enzymes to study eukaryotic DNA methylation. J Mol Biol 118, 27–47. 10.1016/0022-2836(78)90242-5.

13. Luo, G.-Z., Wang, F., Weng, X., Chen, K., Hao, Z., Yu, M., Deng, X., Liu, J., and He, C. (2016). Characterization of eukaryotic DNA N6-methyladenine by a highly sensitive restriction enzyme-assisted sequencing. Nat Commun 7, 11301. 10.1038/ncomms11301.

14. Mahdavi-Amiri, Y., Chung Kim Chung, K., and Hili, R. (2021). Single-nucleotide resolution of N6-adenine methylation sites in DNA and RNA by nitrite sequencing. Chem Sci 12, 606–612. 10.1039/D0SC03509B.

15. Beaulaurier, J., Schadt, E.E., and Fang, G. (2019). Deciphering bacterial epigenomes using modern sequencing technologies. Nat Rev Genet 20, 157–172. 10.1038/s41576-018-0081-3.

16. Schadt, E.E., Banerjee, O., Fang, G., Feng, Z., Wong, W.H., Zhang, X., Kislyuk, A., Clark, T.A., Luong, K., Keren-Paz, A., et al. (2013). Modeling kinetic rate variation in third generation DNA sequencing data to detect putative modifications to DNA bases. Genome Res 23, 129–141. 10.1101/gr.136739.111.

17. Zhu, S., Beaulaurier, J., Deikus, G., Wu, T.P., Strahl, M., Hao, Z., Luo, G., Gregory, J.A., Chess, A., He, C., et al. (2018). Mapping and characterizing N6-methyladenine in eukaryotic genomes using single-molecule real-time sequencing. Genome Res 28, 1067–1078. 10.1101/gr.231068.117.

18. Engel, J.D., and von Hippel, P.H. (1974). Effects of methylation on the stability of nucleic acid conformations. Monomer level. Biochemistry 13, 4143–4158. 10.1021/bi00717a013.

19. Engel, J.D., and von Hippel, P.H. (1978). Effects of methylation on the stability of nucleic acid conformations. Studies at the polymer level. Journal of Biological Chemistry 253, 927–934. 10.1016/S0021-9258(17)38193-0.

20. Harcourt, E.M., Ehrenschwender, T., Batista, P.J., Chang, H.Y., and Kool, E.T. (2013). Identification of a Selective Polymerase Enables Detection of N6-Methyladenosine in RNA. J Am Chem Soc 135, 19079–19082. 10.1021/ja4105792.

21. Sismour, A.M. (2005). The use of thymidine analogs to improve the replication of an extra DNA base pair: a synthetic biological system. Nucleic Acids Res 33, 5640–5646. 10.1093/nar/gki873.

22. Pugliese, K.M., Gul, O.T., Choi, Y., Olsen, T.J., Sims, P.C., Collins, P.G., and Weiss, G.A. (2015). Processive Incorporation of Deoxynucleoside Triphosphate Analogs by Single-Molecule DNA Polymerase I (Klenow Fragment) Nanocircuits. J Am Chem Soc 137, 9587–9594. 10.1021/jacs.5b02074.

23. Koerber, J.T., Maheshri, N., Kaspar, B.K., and Schaffer, D. v (2006). Construction of diverse adeno-associated viral libraries for directed evolution of enhanced gene delivery vehicles. Nat Protoc 1, 701–706. 10.1038/nprot.2006.93.

24. Fromant, M., Blanquet, S., and Plateau, P. (1995). Direct Random Mutagenesis of Gene-Sized DNA Fragments Using Polymerase Chain Reaction. Anal Biochem 224, 347–353. 10.1006/abio.1995.1050.

25. Jia, G., Fu, Y., Zhao, X., Dai, Q., Zheng, G., Yang, Y., Yi, C., Lindahl, T., Pan, T., Yang, Y.-G., et al. (2011). N6-Methyladenosine in nuclear RNA is a major substrate of the obesity-associated FTO. Nat Chem Biol 7, 885–887. 10.1038/nchembio.687.

26. Jia, G., Fu, Y., Zhao, X., Dai, Q., Zheng, G., Yang, Y., Yi, C., Lindahl, T., Pan, T., Yang, Y.-G., et al. (2011). N6-Methyladenosine in nuclear RNA is a major substrate of the obesity-associated FTO. Nat Chem Biol 7, 885–887. 10.1038/nchembio.687.

27. Zhang, X., Wei, L.-H., Wang, Y., Xiao, Y., Liu, J., Zhang, W., Yan, N., Amu, G., Tang, X., Zhang, L., et al. (2019). Structural insights into FTO’s catalytic mechanism for the demethylation of multiple RNA substrates. Proceedings of the National Academy of Sciences 116, 2919–2924. 10.1073/pnas.1820574116.

28. Xie, Q., Wu, T.P., Gimple, R.C., Li, Z., Prager, B.C., Wu, Q., Yu, Y., Wang, P., Wang, Y., Gorkin, D.U., et al. (2018). N-methyladenine DNA Modification in Glioblastoma. Cell 175, 1228–1243.e20. 10.1016/j.cell.2018.10.006.

29. Zhang, M., Yang, S., Nelakanti, R., Zhao, W., Liu, G., Li, Z., Liu, X., Wu, T., Xiao, A., and Li, H. (2020). Mammalian ALKBH1 serves as an N6-mA demethylase of unpairing DNA. Cell Res 30, 197–210. 10.1038/s41422-019-0237-5.

30. Beaulaurier, J., Zhang, X.-S., Zhu, S., Sebra, R., Rosenbluh, C., Deikus, G., Shen, N., Munera, D., Waldor, M.K., Chess, A., et al. (2015). Single molecule-level detection and long read-based phasing of epigenetic variations in bacterial methylomes. Nat Commun 6, 7438. 10.1038/ncomms8438.

31. McIntyre, A.B.R., Alexander, N., Grigorev, K., Bezdan, D., Sichtig, H., Chiu, C.Y., and Mason, C.E. (2019). Single-molecule sequencing detection of N6-methyladenine in microbial reference materials. Nat Commun 10, 579. 10.1038/s41467-019-08289-9.

32. Tapella, R., Ashby, M., Sethuraman, A., and Rhall, P.B. Methylome analysis technical note. https://github.com/PacificBiosciences/Bioinformatics-Training/wiki/Methylome-Analysis-Technical-Note.

33. Geier, G.E., and Modrich, P. (1979). Recognition sequence of the dam methylase of Escherichia coli K12 and mode of cleavage of Dpn I endonuclease. J Biol Chem 254, 1408–1413.

34. Powell, L.M., Lejeune, E., Hussain, F.S., Cronshaw, A.D., Kelly, S.M., Price, N.C., and Dryden, D.T.F. (2003). Assembly of EcoKI DNA methyltransferase requires the C-terminal region of the HsdM modification subunit. Biophys Chem 103, 129–137. 10.1016/S0301-4622(02)00251-X.

35. Kurylo, C.M., Alexander, N., Dass, R.A., Parks, M.M., Altman, R.A., Vincent, C.T., Mason, C.E., and Blanchard, S.C. (2016). Genome Sequence and Analysis of *Escherichia coli* MRE600, a Colicinogenic, Nonmotile Strain that Lacks RNase I and the Type I Methyltransferase, EcoKI. Genome Biol Evol 8, 742–752. 10.1093/gbe/evw008.

36. Mustafa, M.F., Fakurazi, S., Abdullah, M.A., and Maniam, S. (2020). Pathogenic Mitochondria DNA Mutations: Current Detection Tools and Interventions. Genes (Basel) 11, 192. 10.3390/genes11020192.

37. Chen, L.-Q., Zhang, Z., Chen, H.-X., Xi, J.-F., Liu, X.-H., Ma, D.-Z., Zhong, Y.-H., Ng, W.H., Chen, T., Mak, D.W., et al. (2022). High-precision mapping reveals rare N6-deoxyadenosine methylation in the mammalian genome. Cell Discov 8, 138. 10.1038/s41421-022-00484-1.

38. Marinov, G.K., Wang, Y.E., Chan, D., and Wold, B.J. (2014). Evidence for Site-Specific Occupancy of the Mitochondrial Genome by Nuclear Transcription Factors. PLoS One 9, e84713. 10.1371/journal.pone.0084713.

39. Li, Z., Zhao, S., Nelakanti, R. v., Lin, K., Wu, T.P., Alderman, M.H., Guo, C., Wang, P., Zhang, M., Min, W., et al. (2020). N6-methyladenine in DNA antagonizes SATB1 in early development. Nature 583, 625–630. 10.1038/s41586-020-2500-9.

40. Liu, J., Zhu, Y., Luo, G.-Z., Wang, X., Yue, Y., Wang, X., Zong, X., Chen, K., Yin, H., Fu, Y., et al. (2016). Abundant DNA 6mA methylation during early embryogenesis of zebrafish and pig. Nat Commun 7, 13052. 10.1038/ncomms13052.

41. Wu, T.P., Wang, T., Seetin, M.G., Lai, Y., Zhu, S., Lin, K., Liu, Y., Byrum, S.D., Mackintosh, S.G., Zhong, M., et al. (2016). DNA methylation on N6-adenine in mammalian embryonic stem cells. Nature 532, 329–333. 10.1038/nature17640.

42. Li, X., Zhao, Q., Wei, W., Lin, Q., Magnan, C., Emami, M.R., Wearick-Silva, L.E., Viola, T.W., Marshall, P.R., Yin, J., et al. (2019). The DNA modification N6-methyl-2’-deoxyadenosine (m6dA) drives activity-induced gene expression and is required for fear extinction. Nat Neurosci 22, 534–544. 10.1038/s41593-019-0339-x.

43. Kweon, S.-M., Chen, Y., Moon, E., Kvederaviciutė, K., Klimasauskas, S., and Feldman, D.E. (2019). An Adversarial DNA N6-Methyladenine-Sensor Network Preserves Polycomb Silencing. Mol Cell 74, 1138–1147.e6. 10.1016/j.molcel.2019.03.018.

44. Xiao, C.-L., Zhu, S., He, M., Chen, D., Zhang, Q., Chen, Y., Yu, G., Liu, J., Xie, S.-Q., Luo, F., et al. (2018). N6-Methyladenine DNA Modification in the Human Genome. Mol Cell 71, 306–318.e7. 10.1016/j.molcel.2018.06.015.

45. Yao, B., Cheng, Y., Wang, Z., Li, Y., Chen, L., Huang, L., Zhang, W., Chen, D., Wu, H., Tang, B., et al. (2017). DNA N6-methyladenine is dynamically regulated in the mouse brain following environmental stress. Nat Commun 8, 1122. 10.1038/s41467-017-01195-y.

46. Suen, T.C., and Hung, M.C. (1991). c-myc reverses neu-induced transformed morphology by transcriptional repression. Mol Cell Biol 11, 354–362. 10.1128/mcb.11.1.354-362.1991.

47. Schechter, A.L., Stern, D.F., Vaidyanathan, L., Decker, S.J., Drebin, J.A., Greene, M.I., and Weinberg, R.A. (1984). The neu oncogene: an erb-B-related gene encoding a 185,000-Mr tumour antigen. Nature 312, 513–516. 10.1038/312513a0.

48. Drebin, J.A., Stern, D.F., Link, V.C., Weinberg, R.A., and Greene, M.I. (1984). Monoclonal antibodies identify a cell-surface antigen associated with an activated cellular oncogene. Nature 312, 545–548. 10.1038/312545a0.

49. Wang, Y., Huang, N., Li, H., Liu, S., Chen, X., Yu, S., Wu, N., Bian, X.-W., Shen, H.-Y., Li, C., et al. (2017). Promoting oligodendroglial-oriented differentiation of glioma stem cell: a repurposing of quetiapine for the treatment of malignant glioma. Oncotarget 8, 37511–37524. 10.18632/oncotarget.16400.

50. Reindl, J., Shevtsov, M., Dollinger, G., Stangl, S., and Multhoff, G. (2019). Membrane Hsp70-supported cell-to-cell connections via tunneling nanotubes revealed by live-cell STED nanoscopy. Cell Stress Chaperones 24, 213–221. 10.1007/s12192-018-00958-w.

51. Wheeler, M.A., Jaronen, M., Covacu, R., Zandee, S.E.J., Scalisi, G., Rothhammer, V., Tjon, E.C., Chao, C.-C., Kenison, J.E., Blain, M., et al. (2019). Environmental Control of Astrocyte Pathogenic Activities in CNS Inflammation. Cell 176, 581–596.e18. 10.1016/j.cell.2018.12.012.

52. Wan, Q., Ni, L., Wu, L., Zhang, L., Liu, M., and Jiang, X. (2015). The determination of sex type of the cultured murine cell with quantitative PCR technique. Hum Cell 28, 154– 157. 10.1007/s13577-015-0109-3.

53. Raccaud, M., Friman, E.T., Alber, A.B., Agarwal, H., Deluz, C., Kuhn, T., Gebhardt, J.C.M., and Suter, D.M. (2019). Mitotic chromosome binding predicts transcription factor properties in interphase. Nat Commun 10, 487. 10.1038/s41467-019-08417-5.

54. Bayik, D., Bartels, C.F., Lovrenert, K., Watson, D.C., Zhang, D., Kay, K., Lee, J., Lauko, A., Johnson, S., Lo, A., et al. (2022). Distinct cell adhesion signature defines glioblastoma myeloid-derived suppressor cell subsets. Cancer Res. 10.1158/0008-5472.CAN-21-3840.

55. Liu, S., Yin, F., Fan, W., Wang, S., Guo, X., Zhang, J., Tian, Z., and Fan, M. (2012). Over-expression of BMPR-IB reduces the malignancy of glioblastoma cells by upregulation of p21 and p27Kip1. Journal of Experimental & Clinical Cancer Research 31, 52. 10.1186/1756-9966-31-52.

56. Lee, J., Son, M.J., Woolard, K., Donin, N.M., Li, A., Cheng, C.H., Kotliarova, S., Kotliarov, Y., Walling, J., Ahn, S., et al. (2008). Epigenetic-Mediated Dysfunction of the Bone Morphogenetic Protein Pathway Inhibits Differentiation of Glioblastoma-Initiating Cells. Cancer Cell 13, 69–80. 10.1016/j.ccr.2007.12.005.

57. Hong, T., Yuan, Y., Wang, T., Ma, J., Yao, Q., Hua, X., Xia, Y., and Zhou, X. (2017). Selective detection of N6-methyladenine in DNA via metal ion-mediated replication and rolling circle amplification. Chem Sci 8, 200–205. 10.1039/C6SC02271E.

58. DUNN, D.B., and SMITH, J.D. (1955). Occurrence of a New Base in the Deoxyribonucleic Acid of a Strain of Bacterium Coli. Nature 175, 336–337. 10.1038/175336a0.

59. Beh, L.Y., Debelouchina, G.T., Clay, D.M., Thompson, R.E., Lindblad, K.A., Hutton, E.R., Bracht, J.R., Sebra, R.P., Muir, T.W., and Landweber, L.F. (2019). Identification of a DNA N6-Adenine Methyltransferase Complex and Its Impact on Chromatin Organization. Cell 177, 1781–1796.e25. 10.1016/j.cell.2019.04.028.

60. Mondo, S.J., Dannebaum, R.O., Kuo, R.C., Louie, K.B., Bewick, A.J., LaButti, K., Haridas, S., Kuo, A., Salamov, A., Ahrendt, S.R., et al. (2017). Widespread adenine N6-methylation of active genes in fungi. Nat Genet 49, 964–968. 10.1038/ng.3859.

61. He, S., Zhang, G., Wang, J., Gao, Y., Sun, R., Cao, Z., Chen, Z., Zheng, X., Yuan, J., Luo, Y., et al. (2019). 6mA-DNA-binding factor Jumu controls maternal-to-zygotic transition upstream of Zelda. Nat Commun 10, 2219. 10.1038/s41467-019-10202-3.

62. Zhang, G., Huang, H., Liu, D., Cheng, Y., Liu, X., Zhang, W., Yin, R., Zhang, D., Zhang, P., Liu, J., et al. (2015). N6-Methyladenine DNA Modification in Drosophila. Cell 161, 893–906. 10.1016/j.cell.2015.04.018.

63. Greer, E.L., Blanco, M.A., Gu, L., Sendinc, E., Liu, J., Aristizábal-Corrales, D., Hsu, C.-H., Aravind, L., He, C., and Shi, Y. (2015). DNA Methylation on N6-Adenine in C. elegans. Cell 161, 868–878. 10.1016/j.cell.2015.04.005.

64. Luo, G.-Z., and He, C. (2017). DNA N6-methyladenine in metazoans: functional epigenetic mark or bystander? Nat Struct Mol Biol 24, 503–506. 10.1038/nsmb.3412.

65. Liu, J., Yue, Y., and He, C. (2015). Preparation of Human Nuclear RNA m6A Methyltransferases and Demethylases and Biochemical Characterization of Their Catalytic Activity. In, pp. 117–130. 10.1016/bs.mie.2015.03.013.

66. Raine, A., Manlig, E., Wahlberg, P., Syvänen, A.-C., and Nordlund, J. (2017). SPlinted Ligation Adapter Tagging (SPLAT), a novel library preparation method for whole genome bisulphite sequencing. Nucleic Acids Res 45, e36–e36. 10.1093/nar/gkw1110.

67. Tunster, S.J. (2017). Genetic sex determination of mice by simplex PCR. Biol Sex Differ 8, 31. 10.1186/s13293-017-0154-6.

